# Implications of differential size-scaling of cell-cycle regulators on cell size homeostasis

**DOI:** 10.1101/2022.11.30.518453

**Authors:** Xiangrui Ji, Jie Lin

**Affiliations:** Yuanpei College, Peking University, Beijing, China; Center for Quantitative Biology, Academy for Advanced Interdisciplinary Studies, Peking University, Beijing, China; Peking-Tsinghua Center for Life Sciences, Academy for Advanced Interdisciplinary Studies, Peking University, Beijing, China

## Abstract

Accurate timing of division and size homeostasis is crucial for cells. A potential mechanism for cells to decide the timing of division is the differential scaling of regulatory protein copy numbers with cell size. However, it remains unclear whether such a mechanism can lead to robust growth and division, and how the scaling behaviors of regulatory proteins influence the cell size distribution. Here we study a mathematical model combining gene expression and cell growth, in which the cell-cycle activators scale superlinearly with cell size while the inhibitors scale sublinearly. The cell divides once the ratio of their concentrations reaches a threshold value. We find that the cell can robustly grow and divide within a finite range of the threshold value with the cell size proportional to the ploidy. In a stochastic version of the model, the cell size at division is uncorrelated with that at birth. Also, the more differential the cell-size scaling of the cell-cycle regulators is, the narrower the cell-size distribution is. Intriguingly, our model with multiple regulators rationalizes the observation that after the deletion of a single regulator, the coefficient of variation of cell size remains roughly the same though the average cell size changes significantly. Our work reveals that the differential scaling of cell-cycle regulators provides a robust mechanism of cell size control.

**Author summary:** How cells determine the timing of cell division is a fundamental question of cell biology. It has been found that the concentration of cell-cycle activators tends to increase with cell size, while the concentration of inhibitors tends to decrease. Therefore, an attractive hypothesis is that the ratio of activators to inhibitors may trigger cell division. To investigate this hypothesis quantitatively, we study a model including gene expression and cell growth simultaneously. The cell divides once the activator-to-inhibitor ratio reaches a threshold. Combining theories and simulations, we analyze the conditions of robust cell cycle and the cell size distribution. Our model successfully rationalizes several experimental observations, including the relation between cell size and ploidy, the sizer behavior of cell size control, and the change of the mean and breadth of cell size distribution after regulator deletion.

## Introduction

Cells must coordinate growth and division to keep their sizes within a finite range, which is vital for various biological functions and cellular fitness. For example, a very large cell volume may remodel the proteome and trigger senescence [1–3]. On the other hand, a very small cell volume may also reduce cellular fitness due to the limited protein copy numbers and the resulting large fluctuation in protein concentrations [4–6]. Therefore, cells must evolve ways to measure their sizes to accurately decide the timing of division.

Nearly half a century ago, the seminal work by Fantes et al. [7] already systematically discussed the possible mechanisms for cell size control, though the molecular basis of cell-cycle regulation was largely unclear at that time. Fantes et al. described two threshold types for cell-cycle progression. In one case, cell division is triggered once the number of one activator reaches a threshold. Despite its constant concentration, this activator may form a mitotic structure or titrate the genome to promote cell division. It has been proposed that FtsZ and DnaA in bacteria act through this mechanism [8]. In the other case, the regulator concentration matters. This hypothesis is supported by the observations of different eukaryotes across yeast [9–14], mammals [2, 15], and plants [16, 17] that the concentrations of many cell-cycle regulatory proteins change as the cell size increases.

Recently, Chen et al. proposed that the timing of cell-cycle entry can be achieved through the differential scaling of multiple cell-cycle regulators. If the activators exhibit superlinear scaling with cell size while the inhibitors exhibit sublinear scaling, then at some sufficiently large cell size, the activators dominate over the inhibitors and trigger cell-cycle entry. RNA sequencing also showed that the cell-cycle activators (inhibitors) are enriched in the subset of genes that exhibit increasing (decreasing) mRNA concentrations as the cell size increases [12]. Importantly, this mechanism may shed light on a long-standing mystery, especially for budding yeast: cells with some cell-cycle regulators knocked out can still maintain a narrow cell size distribution [18]. While the mechanism of cell size control based on the differential scaling of cell-cycle regulators is plausible, it is unclear whether it leads to robust growth and division. More quantitatively, how does the differential scaling of regulators determine the cell size distribution? The answers to these questions can deepen our understanding of cell-cycle regulation and cell size control.

Regarding the mechanisms of differential scaling, one of us has recently proposed a gene expression model at the whole-cell level in which the promoters of all genes compete for the resource of RNA polymerases (RNAPs) [19]. The model’s key assumption is that RNAP is rate-limiting for transcription, supported by observations including the direct correlation among RNAP subunit level, RNAP occupancy, and transcription rate [20–22], haploinsufficiency of RNAP subunit genes [23, 24], and increased growth rate after overexpression of RNAP subunits [25]. The model predicts that the effective binding affinities of promoters to RNAPs determine the nonlinear scaling of protein numbers with cell size. A gene with a relatively strong promoter exhibits a decreasing protein concentration as the cell size increases, while a gene with a relatively weak promoter exhibits the opposite behavior. The model suggests that the nonlinear scaling between protein numbers and cell size is an inevitable consequence of heterogeneous promoter strengths in the genome. The relation between the promoter strength and nonlinear scaling of protein level raises the possibility that cell-cycle inhibitors may have strong promoters and therefore exhibit sublinear scaling with cell size, while activators may have weak promoters and exhibit superlinear scaling; thus, the ratio of their concentrations depends on cell size and allows a cell to determine its size and decide when to divide [12, 19].

In the main text, we focus on the case where most proteins are nondegradable while the cell-cycle regulators are degradable, supported by the observations that most proteins are stable in yeast and the minority of short-lived proteins are mainly enriched in the cell cycle category [26]. Our conclusions are qualitatively similar for nondegradable cell-cycle inhibitors (S1 Appendix,S7 Fig). We mainly consider a simple scenario with constant gene copy numbers, rationalized by imagining a cell that replicates its genes right before cell division. Therefore, the gene copy number is constant throughout the cell cycle. We also discuss a modified model in which the cell replicates its genes when the ratio of the activator to the inhibitor rises to a threshold and divides after a constant time (S1 Appendix,S6 Fig). Our conclusions are qualitatively similar between the simple scenario and the modified model. We mainly consider symmetric division for simplicity, but our model equally applies to asymmetric division (S1 Appendix,S11 Fig).

In the following, we first discuss the case of a single activator and a single inhibitor. When the ratio of the activator’s concentration to the inhibitor’s concentration rises to a threshold value *θ*, the cell divides (Fig 1A). Based on the deterministic and stochastic versions of this simplified model, we derive the conditions of robust cell cycle and the expression for the cell size and its coefficient of variation (CV). We rationalize the proportional relation between cell size and ploidy, and the sizer behavior of cell size control. Later, we generalize our model to the case of multiple activators and inhibitors, and explain the experimental observations of mutants with regulators deleted.

**Fig 1.**
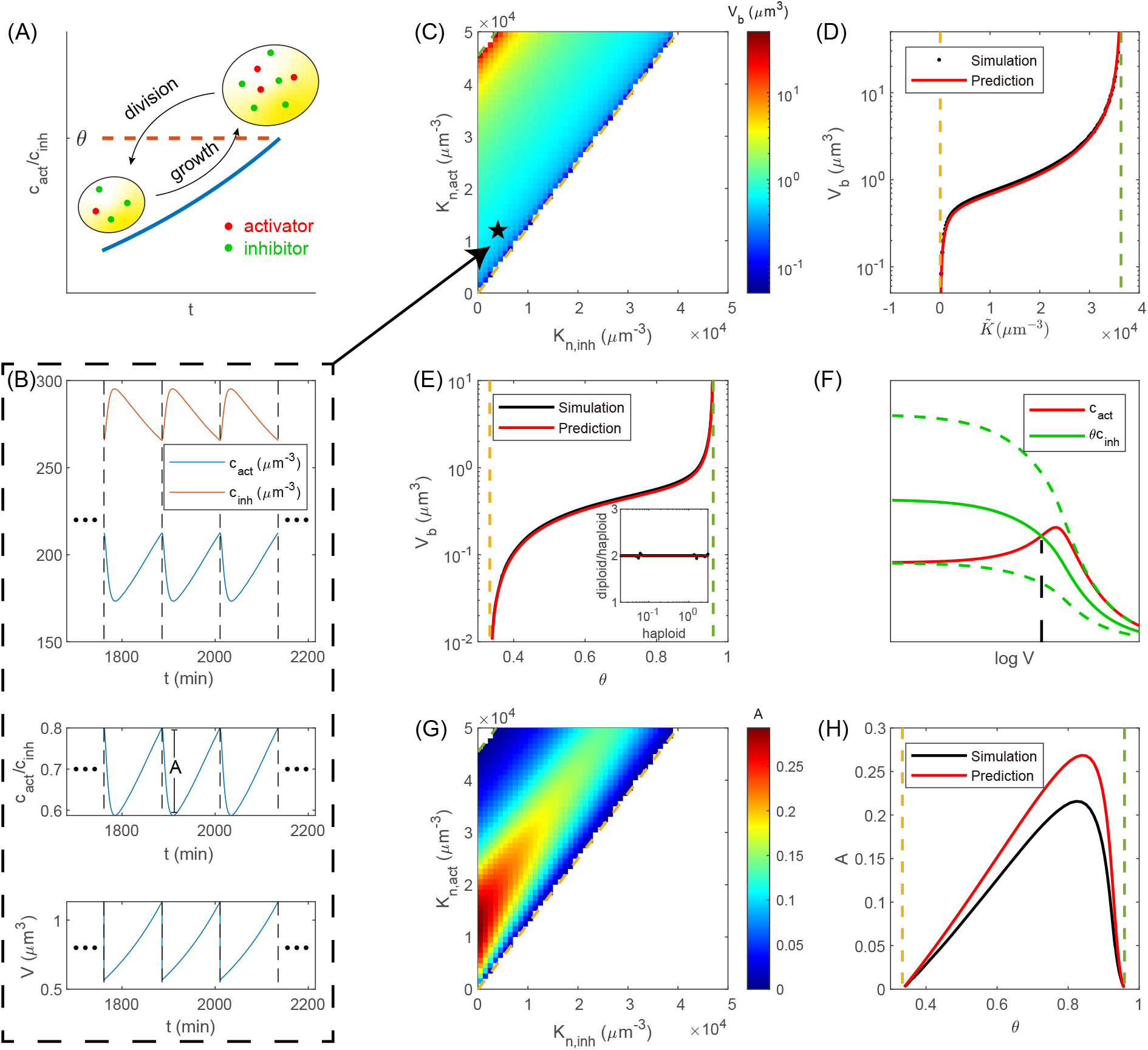
Simulations and theoretical predictions of the cell cycle model. (A) The cell divides symmetrically once the ratio of the activator’s concentration to the inhibitor’s concentration reaches a threshold value. We track one of the daughter cells and simulate a single lineage. (B) (top) The periodic concentrations of the activator *c*_act_ and inhibitor *c*_inh_. The dashed lines mark cell division. The simulation details are in Methods. (middle) The ratio *c*_act_*/c*_inh_ in the periodic steady state. *A* is the amplitude of the periodic oscillation. (bottom) The cell volume *V* also changes periodically. The abrupt drop corresponds to cell division. (C) The heatmap of *V*_*b*_ as a function of *K*_*n*,act_ and *K*_*n*,inh_. The striped color gradient suggests that *V*_*b*_ is a function of a linear combination of *K*_*n*,act_ and *K*_*n*,inh_. As an example, the simulation trajectories for the parameters denoted by the star are shown in (B). (D) Simulations and theoretical predictions of *V*_*b*_ vs. 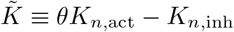. The simulation data here are the same as those in (C). (E) Simulations and theoretical predictions of *V*_*b*_ vs. *θ*. Inset: the ratio between the *V*_*b*_’s of diploid and haploid cells vs. *V*_*b*_ of the haploid cell. The diploid cell size is twice the haploid. Each point corresponds to a given *θ*. (F) Illustration of the origins of the two critical points. The intersection of *c*_act_ and *θc*_inh_ determines the cell size at division *V*_*d*_. Beyond the two critical threshold values (dashed lines), there is no intersection. Between the two critical threshold values, as *θ* increases, the intersection shifts right so that *V*_*d*_ increases with *θ*. (G) The heatmap of *A* as a function of *K*_*n*,act_ and *K*_*n*,inh_. (H) Simulations and theoretical predictions of *A* as a function of *θ*. In (B-D, G), *θ* = 0.8. In (B, E, H), *K*_*n*,act_ = 12000 *µ*m^-3^, *K*_*n*,inh_ = 4000 *µ*m^-3^. The dashed lines in (C-E, G, H) mark the two predicted critical threshold values.

## Results

### Stable cell cycle exists between two critical division thresholds

We use the gene expression model at the whole-cell level which explicitly incorporates the competition between genes for the resource of RNAPs and between mRNAs for the resource of ribosomes [19, 27]. In this model, the concentration of free RNAPs sets the transcription initiation rate through the Michaelis-Menten (MM) mechanism. The MM constant quantifies each gene’s ability to recruit RNAPs. We first consider a model with one activator and one inhibitor and set the MM constants of the activator and inhibitor to be *K*_*n*,act_ and *K*_*n*,inh_ (*K*_*n*,act_ *> K*_*n*,inh_), while the MM constants of other genes are equal to *K*_*n*_ for simplicity. The cell divides once the ratio of the activator’s concentration to the inhibitor’s concentration rises to the threshold value [12, 19],

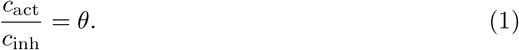

Details of the gene expression model are included in Methods and also in [19].

One of our ultimate goals is to understand how cell size depends on the scaling behaviors of the cell-cycle regulatory proteins. We successfully derive the expression of the cell size at cell birth *V*_*b*_ (see details in Methods). The essence is to find the RNAP number at birth *n*_*b*_. Because the total RNAP concentration *c*_*n*_ is essentially constant, one immediately obtains

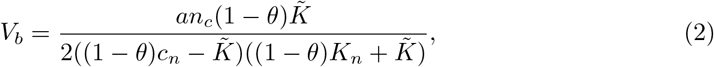

where 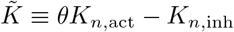. Here *a* is the ratio between the cell volume and the nuclear volume, which is approximately constant, supported by experiments [28, 29]. *n*_*c*_ is the maximum number of RNAPs that the entire genome can hold (Eq 19). Intriguingly, we find that *V*_*b*_ only depends on 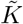, a linear combination of *K*_*n*,act_ and *K*_*n*,inh_ (Fig 1C-D). The comparison between the predictions and simulations shows good agreement (Fig 1D-E, see Methods for the simulation details). We note that the fraction of free RNAPs at cell division *F*_*n,d*_ (see Eq 17 in Methods) must satisfy 0 *< F*_*n,d*_ *<* 1, which is equivalent to requiring the numerator and denominator of Eq 2 to be positive. Therefore, 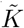 must satisfy

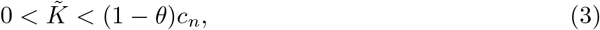

from which we obtain the conditions of a stable cell cycle,

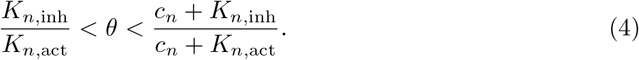

The cell can grow and divide within a finite range of *θ*. The two critical threshold values 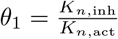 and 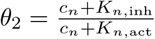 respectively set the lower bound and the upper bound. The predictions of a finite range of *θ* for a stable cell cycle are consistent with simulations (Fig 1B-E).

We provide an intuitive explanation of the critical threshold values in the following. We note that the inhibitor’s concentration keeps decreasing as the cell size increases since its advantage over other proteins becomes weaker and weaker with the increasing concentration of free RNAPs. The activator’s concentration first increases as we predict. However, when the cell size is so big that genes get saturated by RNAPs, and mRNAs get saturated by ribosomes, the protein production rates will also reach their maximum values, which is called Phase 3 of gene expression in [27]. Mathematically, this corresponds to the situation in which the free RNAP concentration is above most genes’ transcription MM constants (Eq 6) and the free ribosome concentration is above most mRNAs’ translation MM constants (Eq 8). Thus, the protein numbers of the degradable activator and inhibitor both become constant, and their concentrations decrease inversely with the cell size in the large cell size limit. Therefore, for the two curves of *c*_act_ and *θc*_inh_ to cross, the threshold *θ* must be within a finite range (Fig 1F).

In the modified model incorporating gene replication (S1 Appendix and S6 Fig), we only need to slightly modify the expression of *V*_*b*_, and the two critical threshold values remain the same. For the case of nondegradable inhibitor, we also derive *V*_*b*_ and provided an illustration for the critical threshold values (S1 Appendix and S7 FigA-B). It is also possible that one mechanism, such as activator-accumulation or inhibitor-dilution, dominates or that the cell integrates the information from each regulator through logic gates [30, 31]. We also explore these cases and perform similar analysis (S1 Appendix and S8 Fig).

We define *A* as the amplitude of the oscillation of *c*_act_*/c*_inh_, i.e., the difference between *θ* and the minimum of *c*_act_*/c*_inh_ in one cell cycle (Fig 1B, middle). We find that the amplitude *A* approaches 0 as *θ* approaches the two critical threshold values (Fig 1G-H). The critical point *θ*_1_ corresponds to *F*_*n,d*_ *→* 0 (see details in Methods). In this limit, the fraction of free RNAPs at cell birth *F*_*n,b*_ also approaches zero since the RNAP number decreases after cell division. The critical point *θ*_2_ corresponds to *F*_*n,d*_ *→* 1 (Methods). In this limit, almost all RNAPs are free, which means that the genes are fully saturated by RNAPs at cell division, and therefore *F*_*n,b*_ is still close to 1. In both cases, *F*_*n,b*_ is close to *F*_*n,d*_. On the other hand, we note that the maximum and minimum of *c*_act_*/c*_inh_ are respectively set by the maximum and minimum of *F*_*n*_ (Eq 22), i.e., *F*_*n,d*_ and *F*_*n,b*_. Therefore, the amplitude *A* approaches 0 as *θ* approaches the two critical threshold values. We derive an approximate expression of *A* (Methods), which agrees well with simulations (Fig 1H). Later we show that the magnitude of *A* is crucial for a stable cell cycle when the threshold *θ* is subject to noise.

### The cell size at birth is proportional to the ploidy

It is well known that the cell size is usually proportional to the ploidy [32–34]. Previous hypotheses about this observation were reviewed in [35, 36]. Our model provides a new perspective on the linear relationship between cell size and ploidy. We remark that our model is invariant as long as the size-to-ploidy ratio is constant. This can be seen from Eq 7 where the doubling of cell size and gene copy numbers leaves the fraction of free RNAPs *F*_*n*_ invariant. Note that the above argument requires that the RNAP concentration *c*_*n*_ is independent of ploidy, which is true in our model (Eq 14) and also supported by experiments [2]. Therefore, the ratio *c*_act_*/c*_inh_ in a diploid cell is the same as that in a haploid cell as long as the size of the diploid cell is twice the size of the haploid cell. Thus, the cell volume at birth *V*_*b*_ doubles in the diploid cell relative to the haploid cell (Fig 1E, inset). This can also be seen from Eq 2, where *n*_*c*_, the total number of RNAPs that can simultaneously bind to the genome, is proportional to ploidy (see Eq 19). In summary, the free RNAP fraction *F*_*n*_ measures the size-to-ploidy ratio and passes on this information to *c*_act_ and *c*_inh_, which then determine the cell size at division. Our model is based on the limiting role of RNAP, supported by experiments [20–25]. Our model predicts that *F*_*n*_ increases with the size-to-ploidy ratio (see Eq 7), consistent with the observation that the number of RNAPs on the genome subscales with cell size [22].

### The cell cycle model with stochastic division threshold reveals a sizer behavior

We extend the model by adding noise to the division threshold *θ* to mimic randomness in real biological systems [4–6]. We assume that the threshold of each generation is independent and identically distributed. Since the threshold value is unlikely to be zero or infinite in real biological systems, we assume *θ* to be within a finite interval 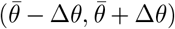 where 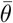 is the mean value. For a given parameter set, we simulate 5000 generations and trace one of the two daughter cells after division (Fig 2A-C). Intriguingly, we find that the cell size at division *V*_*d*_ is uncorrelated with the cell size at birth *V*_*b*_ (Fig 2D), in concert with the sizer behavior found in experiments [37–39]. Therefore, Δ*V ≡ V*_*d*_ *− V*_*b*_ and the doubling time *T*_*D*_ are negatively correlated with *V*_*b*_ (Fig 2E-F). The sizer behavior is robust against the parameter 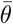 (Fig 3A). This can be seen from Eq 2, where the cell size at cell birth is independent of that of the previous cell cycle. Intuitively, the sizer behavior is due to the degradation of the regulatory proteins: their numbers mainly depend on the current cell size and are insensitive to the history. Sizer due to the fast degradation of cell-cycle regulators is also proposed in [40]. We also study the correlation between *V*_*d*_ and *V*_*b*_ for the case of nondegradable inhibitor and also find an approximately zero correlation between *V*_*d*_ and *V*_*b*_ (S1 Appendix and S7 FigC-E).

**Fig 2.**
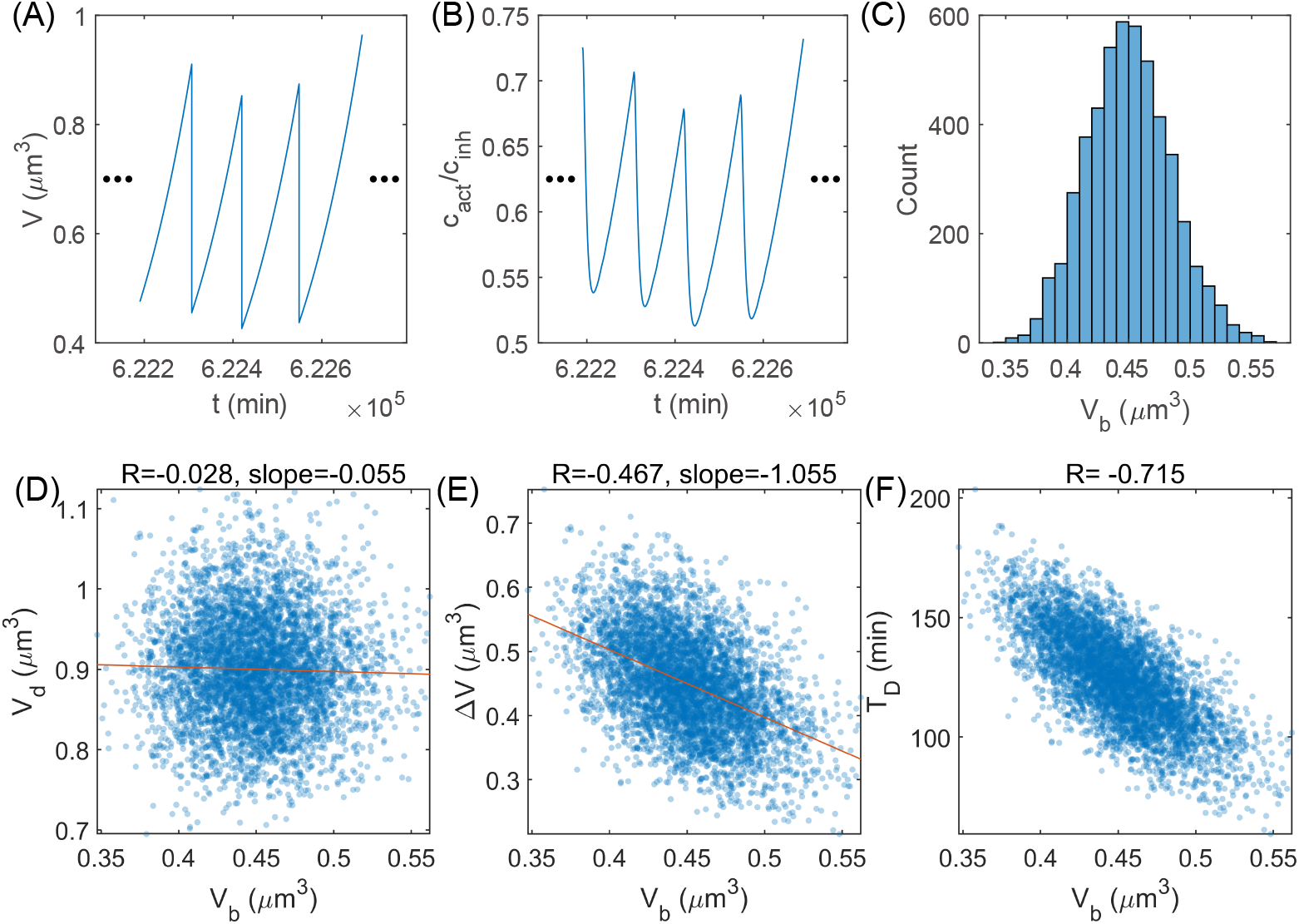
Simulations of the stochastic cell cycle model. (A) The time trajectory of cell size *V* . The cell can grow and divide sustainably. (B) The time trajectory of *c*_act_*/c*_inh_. (C) The distribution of cell size at birth *V*_*b*_. (D) *V*_*d*_ vs. *V*_*b*_. (E) Δ*V* vs. *V*_*b*_. (F) *T*_*D*_ vs. *V*_*b*_. In (D-F), each point represents one cell cycle. The Pearson correlation coefficient *R* and the slope of the linear regression are shown in the title. In all panels, *K*_*n*,act_ = 12000 *µ*m^-3^, *K*_*n*,inh_ = 4000 *µ*m^-3^, *θ* follows a normal distribution *N* (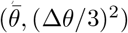, (Δ*θ/*3)^2^), where 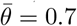, Δ*θ* = 0.1. If *θ* ∉ 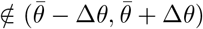, we reset it as 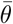.

**Fig 3.**
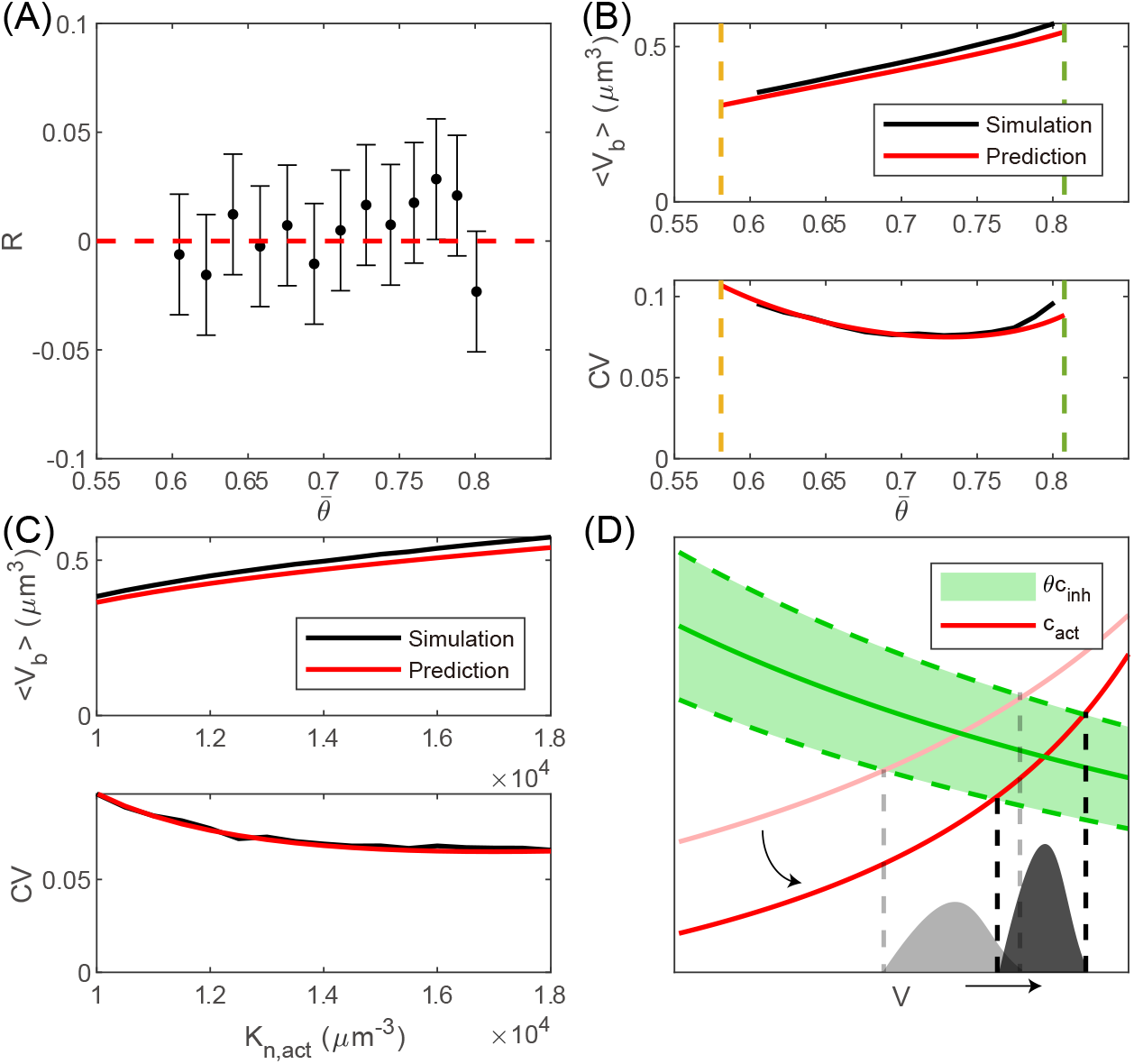
Comparison between theories and simulations of the stochastic model. (A) The Pearson correlation coefficient *R* between *V*_*d*_ and *V*_*b*_ is close to 0 for a range of 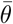. The error bars are the lower and upper bounds for a 95% confidence interval. (B) The average and CV of the cell size at birth *V*_*b*_ for a range of 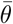. The dashed lines mark the predicted critical threshold values in the presence of noise. In (A-B), *K*_*n*,act_ = 12000 *µ*m^-3^, *K*_*n*,inh_ = 4000 *µ*m^-3^. (C) ⟨*V*_*b*_⟩increases with *K*_*n*,act_, and the CV of *V*_*b*_ decreases with *K*_*n*,act_. *K*_*n*,inh_ = 4000 *µ*m^-3^, 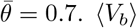. ⟨*V*_*b*_⟩and the CV of *V*_*b*_ as functions of *K*_*n*,inh_ are shown in S2 FigA. In (A-C), Δ*θ* = 0.1. (D) The intersection of *c*_act_ and 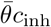 (solid green line) determines the mean cell size at division ⟨*V*_*d*_⟩. The intersections of *c*_act_ with 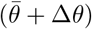 and 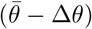 (green dashed lines) set the range of *V*_*d*_. If *K*_*n*,act_ increases, the number of activator becomes more superlinear as a function of cell size, and *c*_act_ shifts to the red line with lower transparency. Therefore, ⟨*V*_*d*_⟩increases while the CV of *V*_*d*_ decreases. Since *V*_*b*_ is half of the *V*_*d*_ of the previous cell cycle, we immediately obtain the distribution of *V*_*b*_. A similar schematic when *K*_*n*,inh_ changes is shown in S2 FigB.

Eukaryotic cells such as the budding yeast *Saccharomyces cerevisiae* [37, 41], the fission yeast *Schizosaccharomyces pombe* [42–44], and mammalian cells [39, 45] are usually imperfect sizers. For budding yeast, this phenomenon could result from the unequal partitioning of Whi5 during asymmetric cell division [46], but it cannot explain the imperfect sizer observed in symmetrically dividing cells. To explain the imperfect sizer, we modify our model by assuming that the cell-cycle entry is probabilistic, with its probability increasing with the activator-to-inhibitor ratio, similar to some previous works [14, 41, 47–49]. This modified model successfully reproduces the imperfect sizer (S1 Appendix,S9 Fig). In addition, we remark that different modes of cell size control in the different cell-cycle phases also contribute to the imperfect sizer over the entire cell cycle [41, 42, 46].

### Differential scaling behaviors of the activator and inhibitor narrow cell size distribution

We also derive the analytical expressions of the mean cell size at birth and the CV of cell size at birth (see details in Methods). The predictions agree well with the simulations under various 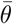, *K*_*n*,act_ and *K*_*n*,inh_ (Fig 3B-C, S2 FigA). We note that when *K*_*n*,act_ increases or *K*_*n*,inh_ decreases, the ⟨*V*_*b*_⟩increases while the CV of *V*_*b*_ decreases, which can be intuitively understood from the cell-size dependence of *c*_act_ and *c*_inh_ (Fig 3D and S2 FigB). Since *K*_*n*,act_ and *K*_*n*,inh_ determine how nonlinear the activator and inhibitor are relative to the cell size, this observation suggests that the more differential the scaling behaviors of the cell-cycle regulators are, the smaller the CV of the cell size distribution is.

In the presence of noise, the range of threshold that allows robust cell growth and division becomes smaller. We find that the mean threshold 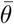 must satisfy (Methods)

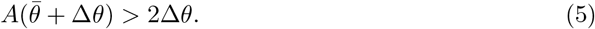

When Δ*θ* = 0, Eq 5 becomes *A*(*θ*) *>* 0, which is precisely the condition of robust division for the deterministic model. We note that ⟨*V*_*b*_⟩can not approach zero or infinity in the presence of noise (Fig 3B), in contrast to the deterministic model. This result suggests that too small or too large cell size is unstable against noise, and cell size must stay in an appropriate range.

### Multiple regulators render robust cell size distribution against gene deletion

Experimentally, it has been found that deleting one regulator’s gene of budding yeast can change the mean cell size significantly, but the CV of cell size is approximately the same before and after deletion [12, 50] (S3 Fig). This suggests that there should be multiple cell-cycle regulators working in concert. To explore the influence of gene deletion, we extend the model to multiple activators and inhibitors. Each regulator has one gene copy, and the numbers of activators and inhibitors are respectively *g*_act_ and *g*_inh_. We assume that all the activators’ promoters have the same effective binding affinity to RNAPs, i.e., *K*_*n*,act_, and all the inhibitors’ promoters have the same effective binding affinity to RNAPs, i.e., *K*_*n*,inh_. The cell divides once the ratio between the concentration of all activators and the concentration of all inhibitors reaches the threshold *θ*, i.e., 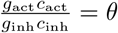, where *c*_act_ (*c*_inh_) is the concentration of a particular activator (inhibitor). We define the equivalent threshold as 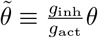 such that the division condition becomes 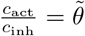. Therefore, all the results above hold as long as we substitute *θ* with 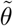 in the deterministic model and substitute 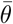 and Δ*θ* with 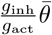 and 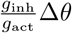 in the stochastic model.

After deleting one activator, the equivalent threshold changes to 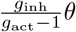, larger than the wild type (WT) value 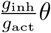. Because *V*_*b*_ increases with *θ* (Fig 1E), the *V*_*b*_ of *activator* Δ is larger than WT, verified by simulations (S4 FigA). In the model of stochastic *θ*, ⟨*V*_*b*_⟩ is also larger after the deletion of one activator (S4 FigB), consistent with experiments [12, 50, 51]. A similar analysis applies to inhibitor deletion. The cell volume becomes smaller after inhibitor deletion (S4 FigA-B). We find that there exists a large parameter range for 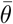 in which the CV’s of *activator* Δ and *inhibitor* Δ are very close to WT (S4 FigB-D), consistent with experiments [12, 50]. For the case of nondegradable inhibitors, the results are qualitatively the same (S10 FigA-D). In S4 FigE-F, we demonstrate schematically the change in cell size distribution. The redundancy of multiple regulators makes the width of cell size distribution insensitive to the deletion of one cell-cycle regulator.

In the following, we provide a more intuitive explanation for the observed constant CV after regulator deletion. We note a wide range of *θ* in which *V*_*b*_ scales linearly with *θ* (Fig 4A). In this range, because *V*_*b*_ *∝ θ*, the CV of *V*_*b*_ is equal to the CV of *θ* in the stochastic model, which is constant. Since deleting the cell-cycle regulator is equivalent to shifting the mean of *θ* while not altering the CV of *θ*, the CV of *V*_*b*_ is robust against gene deletion (Fig 4B).

**Fig 4.**
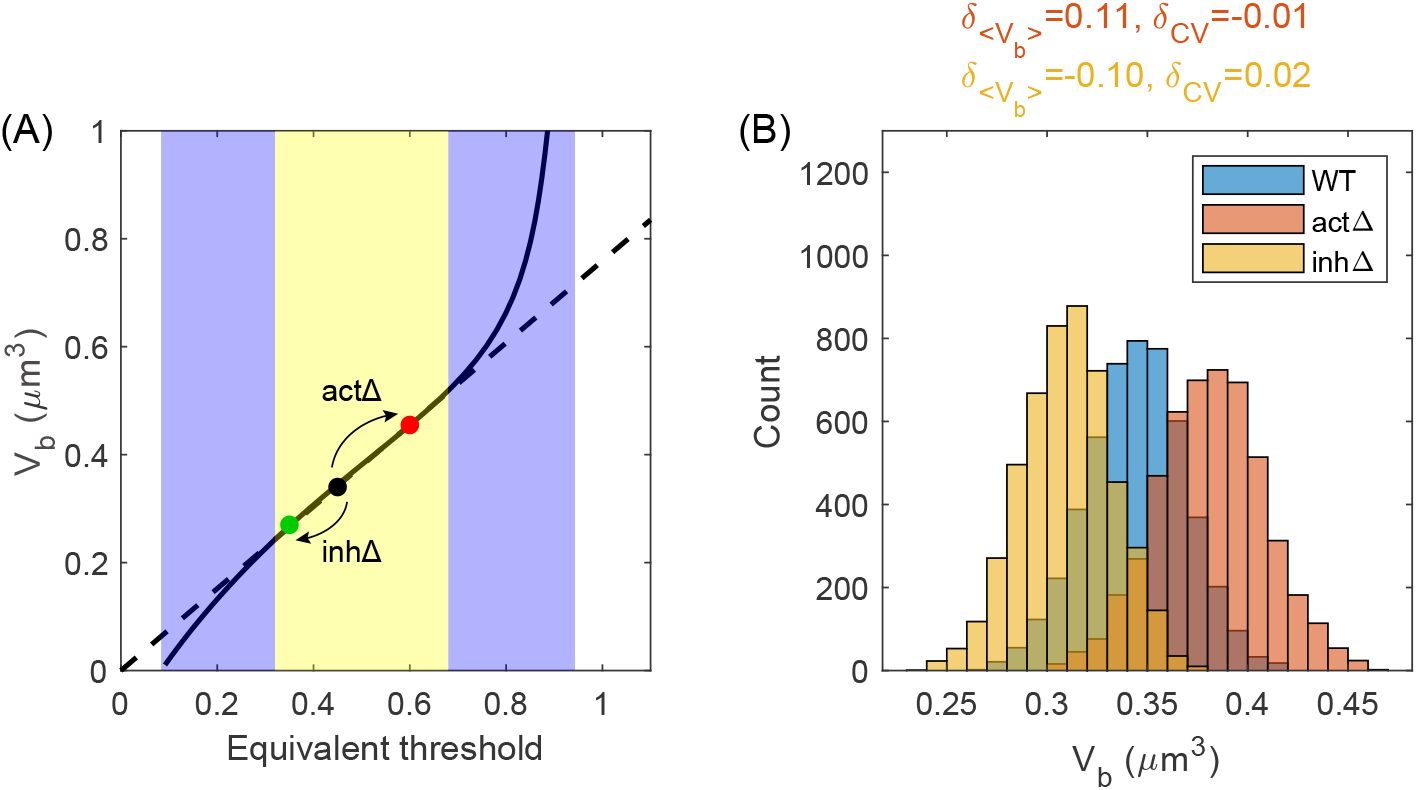
The CV of cell size remains the same after the deletion of one regulator. (A) In our model, the deletion of one activator is equivalent to an increase in the division threshold *θ*, and vice versa for the inhibitor. If the change in equivalent threshold is small, the cell size at birth *V*_*b*_ is still proportional to the threshold (the yellow region) so that the CV of *V*_*b*_ remains constant. (B) The distributions of *V*_*b*_ for WT, *act* Δ and *inh*Δ. The two mutants’ relative changes in average cell size and CV compared with WT are shown at the top 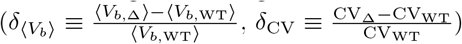. In this figure, *K*_*n*,act_ = 12000 *µ*m^-3^, *K*_*n*,inh_ = 1000 *µ*m^-3^, *g*_act_ = *g*_inh_ = 10, 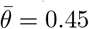, Δ*θ* = 0.1.

Within our model, the simultaneous deletion of multiple activators and inhibitors may lead to a small shift of the equivalent threshold, so the CV in size can change mildly (S12 FigA-B). Meanwhile, our model also predicts that if the shift is drastic and goes beyond the linear regime (e.g., deleting multiple activators *or* multiple inhibitors), the mutant can have a larger CV in size or even become inviable (S12 Fig), in agreement with experiments [12, 44, 52]. Thus, our model provides a unified framework to understand these observations. We acknowledge that our model cannot explain the increase in CV for the fission yeast mutant *wee1-50 cdc25* Δ, which results from the quantized cell-cycle times [43, 53, 54].

We remark that given multiple regulators, the lack of one copy of an inhibitor’s gene in a diploid cell does not necessarily halve the cell size. The redundancy of multiple inhibitors naturally explains the observation that the cell size of the diploid *WHI5* heterozygous mutant is still bigger than the haploid WT cell [9].

## Discussion

In this work, we study a model combining cell size control and the differential scaling of gene expression with cell size. As the cell grows, the ratio between the activator and inhibitor increases and eventually triggers cell division when the ratio is above a threshold value [12, 19]. The activator has a relatively weak promoter and therefore scales superlinearly with cell size. Meanwhile, the inhibitor has a relatively strong promoter and therefore scales sublinearly. Our simplified model captures the essential features of cell size control, and the cell can periodically grow and divide, maintaining size homeostasis. We derive the analytical expression of the cell size and determine the conditions of stable cell cycle, revealing that global gene expression can impose a constraint on cell size control. We also provide a new perspective on the linear relationship between cell size and ploidy [32–34], showing that the free RNAP fraction *F*_*n*_ measures the size-to-ploidy ratio and passes on this information to *c*_act_ and *c*_inh_. All of our theoretical predictions are verified by simulations.

We also propose a stochastic version of our model in which the division threshold is random in each cell cycle. We find that the cell size at division is uncorrelated with the cell size at birth, known as a sizer [37–39]. Intriguingly, we find that the more differential the cell-size scaling of the cell-cycle regulators is, the narrower the cell-size distribution is, which unravels the role of differential scaling in size homeostasis.

We also extend our model to multiple activators and inhibitors. The cell size becomes larger after one activator is deleted, while it becomes smaller after one inhibitor is deleted, which agrees with experiments [12, 50, 51]. Interestingly, while the mean cell size can change significantly after regulator deletion, the CV of the cell size distribution can be approximately constant, consistent with experiments [12, 50]. To our knowledge, no previous theoretical works explained this counter-intuitive phenomenon. Our model further predicts that though the CV of cell size is robust against deletion of one single regulator, it may increase after deletions of multiple activators or multiple inhibitors. It will be interesting to test our prediction by systematically measuring the CV of budding yeast strains with multiple activators or multiple inhibitors knocked out, which is still largely unexplored.

In the experiment that exchanged the promoters of *CLN2* and *WHI5*, the mean and CV of the budding yeast cell size increased [12]. Our model can also reproduce the increase in the mean and CV of cell size after swapping the promoters of the activator and inhibitor (S10 FigE). Meanwhile, we note that the outcome of exchanging *CLN2* and *WHI5* promoters can be more complex due to factors such as regulations acting on the promoters. Whi5 inhibits the transcription of *CLN2* by binding the SBF complex, and Cln2 can phosphorylate Whi5 and dissociate it from SBF, forming a positive feedback loop [55–57]. Therefore, exchanging the *WHI5* promoter and *CLN2* promoter may rewire the network and alter the regulatory relationship. Besides, systematic experiments of exchanging promoters are still lacking, and it is unclear whether the increase in cell size and its CV after promoter exchange is a universal phenomenon. Future experimental work on this issue will be very helpful. Additionally, experiments properly truncating the promoters of the activator and inhibitor might change the binding affinities of promoters without altering the regulatory network, which can help test the prediction that the nonlinear scaling behaviors of regulators influence the breadth of the cell size distribution.

Wang et al. have analyzed the RNA-seq data of [12] and showed that genes exhibiting sublinear scaling tend to have higher mRNA production rates. In particular, negative cell-cycle regulators are enriched in the sublinear regime [19]. It will be helpful to test further our assumption that cell-cycle activators tend to have lower RNAP occupancy due to their weak promoters, and vice versa for inhibitors, e.g., by combining RNA-seq and ChIP-seq data of budding yeast cells arrested in the G1 phase. Meanwhile, we acknowledge that this assumption may not apply to all cell-cycle regulators.

Conceptually, the ideas that limiting RNAP accounts for the increase of transcription rate with cell size [20, 21, 58, 59] and that different recruitment abilities of promoters to RNAP generate nonlinear scaling of gene expression with cell size [19, 46, 60] have been brought forward before. Our work shares some similarities with previous works. Heldt et al. [46] studied a mathematical model of cell size control for budding yeast in which the sublinear scaling of Whi5 is introduced by a strong promoter while Cln3 exhibits linear scaling. We remark that the model by Heldt et al. was designed to understand the cell size control mechanism of budding yeast and, in particular, to explain the size of cells with abnormal *WHI5* copy numbers and rule out alternative mechanisms. However, our model is not constructed for a specific organism and is more poised to obtain some general conclusions that may apply to multiple organisms. Moreover, our gene expression model is built at the whole-cell level and reveals the connection between global gene expression and cell size control. Furthermore, our model with multiple regulators explains the counter-intuitive phenomenon that the CV of cell size can change mildly after the deletion of key regulators [12]. While Heldt et al. assumed a linear size-scaling for activators, we include the possibility of superlinear scaling for activators, given that it could also facilitate cell size homeostasis [12–14]. Other works employing the idea of nonlinear scaling did not include gene expression mechanisms for the nonlinear scaling behaviors therein [61, 62].

Our results support the hypothesis that cells can measure their sizes through the differential scaling of cell-cycle regulatory proteins. Our model is not designed for specific cell-cycle regulators or organisms, so that some specific experimental phenomena could be beyond our model. Nevertheless, our model can be extended by including cell-cycle regulatory networks [52, 63–65].

## Methods

### Details of the gene expression model

We list the variables used in the main text in Table 1. A free RNAP can bind to a promoter and become an initiating RNAP, which can start transcribing or hop off the promoter without transcribing. Thus, the concentration of free RNAPs sets the transcription initiation rate through the Michaelis-Menten (MM) mechanism. The MM constant quantifies each gene’s ability to recruit RNAPs. The time dependence of the mRNA copy number of one particular gene *i* follows [19],

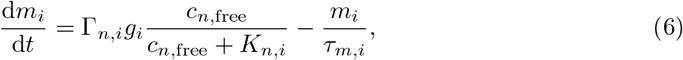

where *c*_*n*,free_ is the free RNAP concentration in the nucleus and *K*_*n,i*_ is the MM constant. A larger *K*_*n,i*_ represents a weaker promoter, while a smaller *K*_*n,i*_ represents a stronger promoter. *g*_*i*_ is the copy number of gene *i* and *τ*_*m,i*_ is the corresponding mRNA lifetime. Γ_*n,i*_ is the rate for a promoter-bound RNAP to become a transcribing RNAP. Interaction between genes due to the limiting resource of RNAPs is through the free RNAP concentration. For one copy of each gene, the number of actively transcribing RNAPs is proportional to the probability that the promoter is bound by an RNAP, based on the assumption that the number of RNAPs starting transcription per unit time is equal to the number of RNAPs finishing transcription per unit time. This assumption is justified by the fact that it usually takes shorter than one minute for an RNAP to finish transcribing a gene [66] so that the system can quickly reach a steady state.

**Table 1.** A summary of the variables.

| Variables | Meaning | Variables | Meaning |
| --- | --- | --- | --- |
| $\Gamma_{n,i}$ | transcription initiation rate of gene $i$ | $\Gamma_{r,i}$ | translation initiation rate of gene $i$ |
| $n$ | RNAP number | $r$ | ribosome number |
| $c_n$ | RNAP concentration | $c_r$ | ribosome concentration |
| $F_n$ | free RNAP fraction | $F_r$ | free ribosome fraction |
| $K_{n,i}$ | transcription MM constant of gene $i$ | $K_{r,i}$ | translation MM constant of gene $i$ |
| $v_n$ | RNAP elongation speed | $v_r$ | ribosome elongation speed |
| $\tau_{m,i}$ | lifetime of mRNAs $i$ | $\tau_{p,i}$ | lifetime of protein $i$ |
| $m_i$ | mRNA number of gene $i$ | $p_i$ | protein number of gene $i$ |
| $g_i$ | gene copy number of gene $i$ | $V$ | cell volume |
| $L_i$ | length of gene $i$ | $a$ | ratio between cell volume and nuclear volume |
| $\rho$ | protein mass per cell volume | $n_c$ | maximum number of RNAPs the entire genome can hold |
| $N$ | total number of genes | $\theta$ | threshold for division |

We introduce the relative fraction of free RNAPs in total RNAPs *F*_*n*_ such that *c*_*n*,free_ = *c*_*n*_*F*_*n*_ where *c*_*n*_ is the concentration of total RNAPs in the nucleus. Using the conservation of the total RNAP number, we can find the free RNAP concentration self-consistently [19],

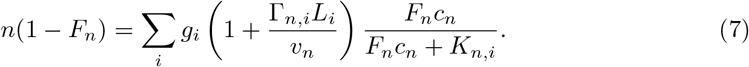

The right side of Eq 7 represents RNAPs binding to the promoters or transcribing. Here, *L*_*i*_ is the length of gene *i* in the unit of codon, and *v*_*n*_ is the RNAP elongation speed. When the number of transcribing RNAPs on one copy of gene *i, n*_*i*_, does not change with time, we must have Γ _*n,i*_*P*_*b,i*_ = *v*_*n*_*n*_*i*_*/L*_*i*_. Here the left side is the number of RNAPs starting transcription per unit time, where *P*_*b,i*_ *≡ F*_*n*_*c*_*n*_*/*(*F*_*n*_*c*_*n*_ + *K*_*n,i*_) is the probability that the promoter is bound by an RNAP. The right side is the number of RNAPs finishing transcribing per unit time. Therefore, we obtain *n*_*i*_ = *P*_*b,i*_ Γ _*n,i*_ *Γ*_*i*_*/v*_*n*_, which is included in the right side of Eq 7.

A similar model as Eq 6 applies to the protein number,

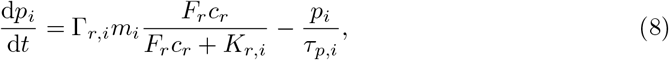

in which the free ribosome concentration sets the initiation rate of translation. In particular, we denote the protein number of RNAPs and ribosomes as *n* and *r*, and set their index as *i* = 1 and *i* = 2 respectively. Γ_*r,i*_ is the translation initiation rate for a ribosome binding on the ribosome-binding site of mRNAs, *c*_*r*_ is the concentration of total ribosomes in the cytoplasm, *K*_*r,i*_ quantifies the binding strength of mRNAs to ribosomes. *τ* _*p,i*_ is the protein lifetime, and *F*_*r*_ is the fraction of free ribosomes in the total pool of ribosomes. *F*_*r*_ satisfies the following equations due to the conservation of total ribosome numbers,

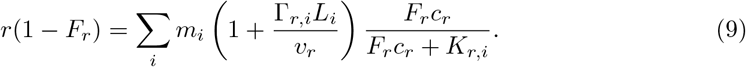

The right side of Eq 9 represents ribosomes binding to the ribosome-binding sites of mRNAs or translating. *v*_*r*_ is the ribosome elongation speed.

In this work, we assume that the cell size is simply proportional to the total protein mass, supported by experiments [67–70]. For simplicity, we assume that all genes except the activator and the inhibitor share the same transcription MM constant *K*_*n*_. Heterogeneous *K*_*n,i*_ among genes has no significant effects on our results (S5 Fig). All genes except those of RNAPs and ribosomes share the same transcription initiation rate Γ_*n*_ (see Details of simulations in Methods). All the mRNAs share the same effective binding strength to ribosomes *K*_*r*_, the same translational initiation rate Γ_*r*_, and the same mRNA lifetime *τ*_*m*_. The cell-cycle regulators share the same protein lifetime *τ*_*p*_ and other proteins are nondegradable.

In the simplified model with a single activator and a single inhibitor, the cell divides once the activator-to-inhibitor ratio *c*_act_*/c*_inh_ rises to the threshold value *θ*. The cell can reach a steady state, growing and dividing periodically (Fig 1B).

### The non-monotonic behavior of *c*_act_*/c*_inh_ in one cell cycle

There is a transient increase in the inhibitor concentration and a transient decrease in the activator concentration right after cell division (Fig 1B). This is because of the reduced RNAP number at cell birth compared to cell division. The fewer total RNAP number leads to fewer free RNAPs given the same gene copy number, which means a decreasing concentration of free RNAPs (S1 Fig). The initial low concentration of free RNAPs lets the inhibitor increase faster than other proteins because of its strong promoter, leading to an increasing inhibitor concentration. However, the advantage of the inhibitor over other proteins becomes weaker as the free RNAP concentration increases since all promoters are fully saturated in the limit of very high free RNAP concentration. Therefore, the inhibitor’s concentration decreases as the cell grows. The opposite situation applies to the activator. After the transient regime, *c*_act_*/c*_inh_ increases, eventually reaches the threshold *θ* and triggers cell division.

A more quantitative explaination is as follows. Using Eq 6 about the activator and the inhibitor, we obtain

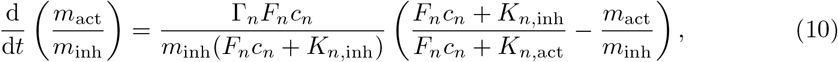

The left-hand side of Eq 7 decreases monotonically with *F*_*n*_, while the right-hand side increases monotonically. Therefore, *F*_*n*_ jumps to a smaller value after cell division since *n* is reduced by half after division. According to Eq 10, the drop in *F*_*n*_ turns 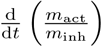 into a negative value, which means that *m*_act_*/m*_inh_ starts to decrease. Consequently, the ratio of the production rates of the activator to that of the inhibitor declines, leading to the reduction of *c*_act_*/c*_inh_. We note that the transient decrease of *c*_act_*/c*_inh_ at the beginning of cell cycle is abrupt in the limit of short lifetimes of mRNAs and regulatory proteins (S1 Appendix). Then, 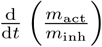 becomes positive again as *F*_*n*_ increases, causing the increase in *c*_act_*/c*_inh_.

### Derivation of the cell size at birth and the two critical threshold values

Assuming a short lifetime of *m*_*i*_ [66] in Eq 6, we can approximate the mRNA copy number as

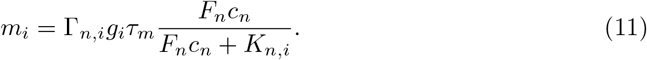

For those proteins that are not cell-cycle regulators, substituting Eq 11 into Eq 8, we get

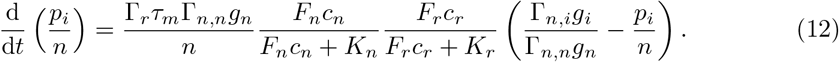

At steady state, 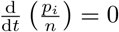. Therefore,

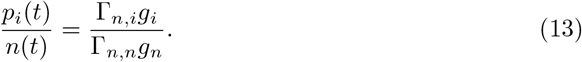

The ratio of copy numbers between two proteins is determined by their gene copy numbers and transcription initiation rates.

The total protein mass of the cell is Σ_*i*_*p*_*i*_*L*_*i*_, in the number of amino acids. The cell size *V* (*t*) = Σ_*i*_ *p*_*i*_(*t*)*L*_*i*_*/ρ*, where *ρ* is the ratio between the total protein mass and cell size, which is a constant in our model, consistent with experiments [67–70]. Therefore, the concentration of total RNAPs in the nucleus is approximately constant and can be approximated as

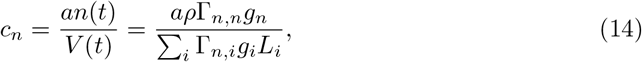

Here we have neglected the contribution of the cell-cycle regulators to the total protein mass. Eq 14 shows that the RNAP concentration is independent of ploidy since the double of all genes’ copy numbers leaves *c*_*n*_ invariant. Assuming a short lifetime *τ*_*p*_, using Eq 11 and the quasi-steady state approximation about *p*_act_ and *p*_inh_ in Eq 8, we get

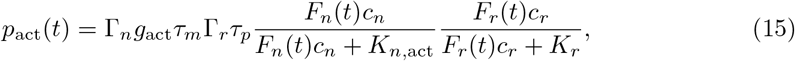

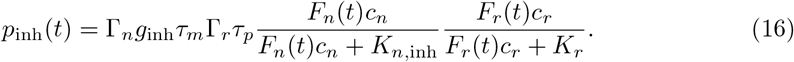

In the simple case of *g*_act_ = *g*_inh_ = 1, substituting Eqs 15, 16 into the division condition Eq 1 (note that *c*_act_*/c*_inh_ = *p*_act_*/p*_inh_), we obtain the fraction of free RNAPs at cell division

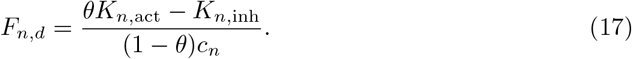

We also simplify Eq 7 using the assumption that *K*_*n,i*_ = *K*_*n*_ for most genes,

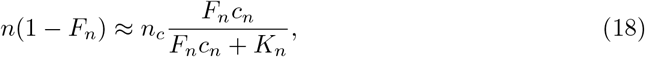

where *n*_*c*_ is the maximum number of RNAPs the entire genome can hold [19],

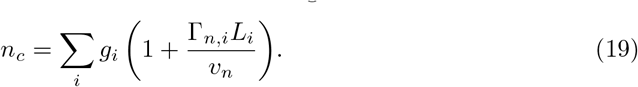

Substituting Eq 17 into Eqs 11,18, we get the mRNA copy number and the RNAP number at cell division

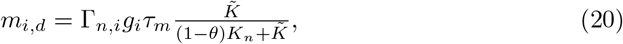

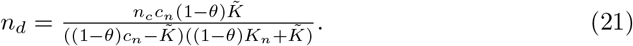

Using *n*_*b*_ = *n*_*d*_*/*2 where *n*_*b*_ is the RNAP number at cell birth and the RNAP concentration Eq 14, we obtain Eq 2 (see S1 Appendix for the discussion of asymmetric division). The condition 0 *< F*_*n,d*_ *<* 1 in Eq 17 leads to the condition of stable cell cycle Eq 3 so that *θ*_1_ corresponds to *F*_*n,d*_ = 0 while *θ*_2_ corresponds to *F*_*n,d*_ = 1. We mathematically prove that the cell cycle is stable against infinitesimal perturbation (S1 Appendix).

### Derivation of the amplitude *A*

Using Eqs 15, 16, the ratio of the activator’s concentration to the inhibitor’s concentration at a given time *t* during the cell cycle can be approximated as

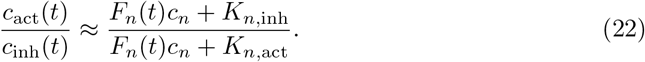

The ratio reaches its minimum when *F*_*n*_ is equal to its minimum value *F*_*n,b*_ at cell birth. Therefore,

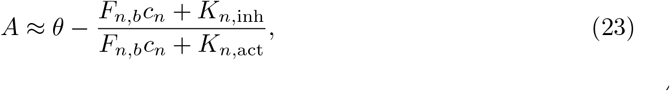

where *F*_*n,b*_ satisfies

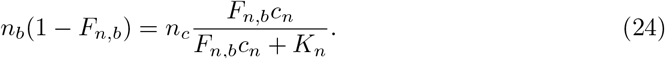

As *θ* increases, *n*_*b*_ increases (Eq 21). According to Eq 24, more RNAPs at cell birth generates more free RNAPs, leading to a larger *F*_*n,b*_. Therefore, 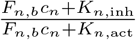 also increases with *θ*, consistent with the fact that *A* first increases and then decreases with *θ*.

### Derivation of ⟨*V*_*b*_ ⟩and the CV of *V*_*b*_

When Δ*θ* is small, a function *f* of *θ* can be approximated as

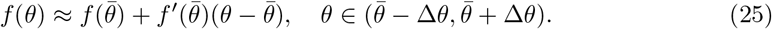

Therefore, we can approximate the average and variance of *f* (*θ*) as

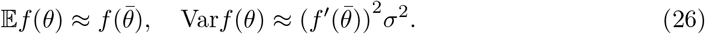

Here *σ*is the standard deviation of *θ*. Using Eqs 2, 26, we obtain

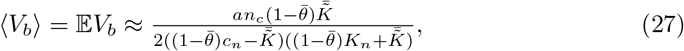

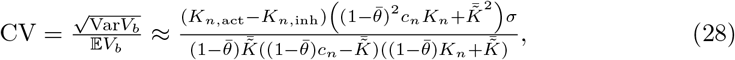

where 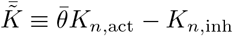.

### The condition of robust cell division for the stochastic model

We denote *θ*_*k*_ as the division threshold value of the *k*-th cell cycle. Given 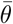 and Δ*θ*, to ensure robust division, we require that the minimum value of the activator-to-inhibitor ratio in the *k*-th cell cycle *θ*_*k−*1_ *− A*(*θ*_*k−*1_) is smaller than *θ*_*k*_, where *A*(*θ*_*k−*1_) is the amplitude of the *k*-th cell cycle, which is a function of *θ*_*k−*1_. This condition must hold for any *θ*_*k−*1_ and *θ*_*k*_. Therefore,

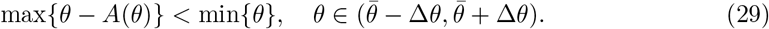

Because *θ − A*(*θ*) increases with *F*_*n,b*_ (see Eq 23) and *F*_*n,b*_ increases with *θ, θ − A*(*θ*) increases with *θ*. Therefore,

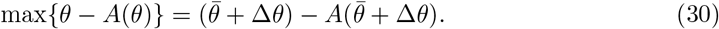

Substituting Eq 30 into Eq 29, we obtain Eq 5.

### Details of simulations

All simulations were done in MATLAB (version R2021b). We summarize some of the parameters used in simulations in Table S1 of S1 Appendix. To find reasonable values of Γ_*n,i*_, we first set *n*_*c*_ = 10^4^ as in [19]. Assuming the number of RNAPs is about 10% of ribosomes i.e. *n* = 0.1*r*, from Eq 13 we get

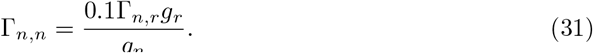

Assuming the length of genes except those of RNAPs and ribosomes is *L*, from Eq 19 we get,

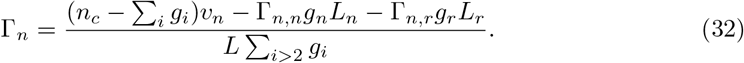

Substituting Eq 11 into Eq 9, we get

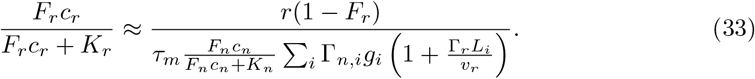

Substituting Eqs 11,33 into Eq 8 for *r*, we find the growth rate as

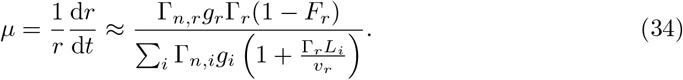

In this work, we set the attempted growth rate *µ* = (ln 2)*/*2 h^-1^. Substituting Eqs 31,32 to Eq 34, and letting *F*_*r*_ = 0, which is merely a numerical convenience, we obtain

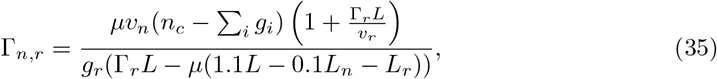

from which we find Γ _*n,n*_ and Γ _*n*_.

Our theoretical results do not rely on the initial values of simulations, but appropriate initial values make it easier to reach the steady state. We first set an attempted cell volume *V*_0_, and then we get *n*(0) = *c*_*n*_*V*_0_*/a* where *c*_*n*_ is calculated using Eq 14. Finally, we determine *p*_*i*_(0) (*i >* 1) using Eq 13 and *m*_*i*_(0) using Eq 11. After symmetric division, mRNAs and proteins are equally distributed to the two daughter cells. We also have a requirement on the minimum cell-cycle duration to avoid cell division immediately after cell birth. Before we take the simulation results, we run the simulations for several generations to ensure that the cell reaches a steady state.

## Supporting information

**S1 Appendix. Supplementary information**.

**S1 Fig. The free RNAP fraction** *F*_*n*_ **drops abruptly at division while the free ribosome fraction** *F*_*r*_ **does not (related to Fig 1)**. *K*_*n*,act_ = 12000 *µ*m^-3^, *K*_*n*,inh_ = 4000 *µ*m^-3^, and *θ* = 0.8.

**S2 Fig. Comparison between theories and simulations of the stochastic model (related to Fig 3)**. (A) ⟨ *V*_*b*_ ⟩ decreases with *K*_*n*,inh_, and the CV of *V*_*b*_ increases with *K*_*n*,inh_. *K*_*n*,act_ = 12000 *µ*m^-3^, 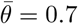, Δ*θ* = 0.1. (B) Illustration of the tendency in (A). The intersection of *c*_inh_ and 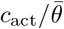 (red solid line) determines ⟨ *V*_*d*_ ⟩. The intersections of *c*_inh_ with 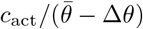 and 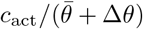 (red dashed lines) set the range of *V*_*d*_. If *K*_*n*,inh_ decreases, the number of inhibitor becomes more sublinear as a function of cell size and *c*_inh_ shifts to the green line with lower transparency. Therefore, ⟨ *V*_*d*_ ⟩ increases while the CV of *V*_*d*_ decreases. Since *V*_*b*_ = *V*_*d*_*/*2, the same conclusions apply to *V*_*b*_.

**S3 Fig. The standard deviation (STD) of cell size vs. the mean cell size at budding of various budding yeast mutants**. Each point represent a budding yeast mutant with one particular gene belonging to the positive or negative regulator category [50] knocked out, and the red line is the linear fit using the equation *y* = *bx*. The *R*^2^ is about 0.92, indicating that the STD is nearly proportional to the mean. Thus, after deleting an activator or inhibitor, the mean cell size at budding can change a lot while the CV of cell size at budding remains roughly the same. The data are from Dataset S3 of [50].

**S4 Fig. Simulations of multiple regulators (related to Fig 4)**. (A) Given a fixed *θ, V*_*b*_ increases after deleting one activator and decreases after deleting one inhibitor. The dashed lines mark the range of *θ* that simultaneously allows WT, *activator* Δ and *inhibitor* Δ to divide. (B) In the stochastic model, ⟨ *V*_*b*_ ⟩ increases after deleting one activator and decreases after deleting one inhibitor. The CV of *V*_*b*_ changes mildly after deleting one regulator. The dashed lines mark the range of 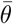 that simultaneously allows WT, *activator* Δ and *inhibitor* Δ to divide. (C) The absolute value of the relative change 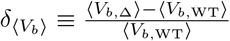 and 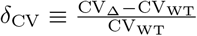 after deleting one activator. (D) 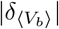 and | *δ*_CV_ | vs. 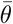 after deleting one inhibitor. The dashed lines in (C-D) have the same meaning as panel (B). There exists a range of viable 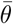 such that the relative change of CV is smaller than that of *(V*_*b*_) after regulator deletion. In (B-D), Δ*θ* = 0.1. In (A-D), *K*_*n*,act_ = 12000 *µ*m^-3^, *K*_*n*,inh_ = 1000 *µ*m^-3^, *g*_act_ = *g*_inh_ = 10. (E) The deletion of one activator shifts the total concentration of all activators (Σ*c*_act_) downward while the total concentration of all inhibitors (Σ*c*_inh_) remains the same. This leads to a change in the mean cell size at division ⟨*V*_*d*_⟩ while the CV of *V*_*d*_ is almost the same. (F) The deletion of one inhibitor shifts the total concentration of all inhibitors (Σ*c*_inh_) downward while the total concentration of all activators (Σ*c*_act_) remain the same. This leads to a change in ⟨*V*_*d*_⟩ while the CV of *V*_*d*_ is almost the same.

**S5 Fig. The model with heterogeneous** *K*_*n,i*_ **among genes**. We simulate a model in which the transcription MM constants are heterogeneous among genes, and compare the simulations with theoretical predictions. (A) *V*_*b*_ vs. 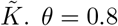. (B) *V*_*b*_ vs. *θ*. (C) *A* vs. *θ*. In (B-C), *K*_*n*,act_ = 12000 *µ*m^-3^, *K*_*n*,inh_ = 4000 *µ*m^-3^. In all panels, the dashed lines mark the two predicted critical threshold values. *K*_*n,i*_ follows a lognormal distribution with the mean *K*_*n*_ = 6000 *µ*m^-3^ and the CV equal to 0.5.

**S6 Fig. The modified model with gene replication**. (A) The concentration of the activator *c*_act_ and inhibitor *c*_inh_, (B) *c*_act_*/c*_inh_ and (C) the cell volume *V* change periodically. (D) The free RNAP fraction *F*_*n*_ drops abruptly at gene replication while the free ribosome fraction *F*_*r*_ does not. In (A-D), the dashed lines mark cell division, and *θ* = 0.7. (E) Simulations and theoretical predictions of *V*_*b*_ vs. *θ*. In this figure, *K*_*n*,act_ = 12000 *µ*m^-3^, *K*_*n*,inh_ = 4000 *µ*m^-3^, *T*_*M*_ = 60 min.

**S7 Fig. The model of nondegradable inhibitor**. (A) Simulations and theoretical predictions of *V*_*b*_ vs. *θ* in the deterministic model. The dashed line marks the predicted critical points *θ*_1_. (B) Illustration of two critical division thresholds. Note that *c*_act_ and *θc*_inh_ may have two intersections. Only the intersection on the left determines *V*_*d*_, because as the cell volume increases, *c*_act_*/c*_inh_ must rise to *θ*. Beyond two critical points (dashed lines), there is no intersection. (C) *V*_*d*_ vs. *V*_*b*_ in the stochastic model. (D) Δ*V* vs. *V*_*b*_. (E) *T*_*D*_ vs. *V*_*b*_. In (C-E), we simulate the stochastic model and each point represents one cell cycle. The Pearson correlation coefficient *R* and slope of the linear regression are shown in the title. 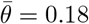, Δ*θ* = 0.018. In (A, C-E), *K*_*n*,act_ = 12000 *µ*m^-3^, *K*_*n*,inh_ = 4000 *µ*m^-3^, *α* = 10.

**S8 Fig. Alternative criteria for cell division**. (A) In this modified model, the cell divides once the activator concentration *c*_act_ increases to the threshold *θ*, and the activator is degradable. (B-C) The cell divides once the inhibitor concentration *c*_inh_ decreases to the threshold *θ*. The inhibitor is degradable in (B) and nondegradable in (C). In each panel, on the left is *c*_act_ (*c*_inh_) vs. *V*, illustrating the origins of the critical points. The intersection of *c*_act_ (*c*_act_) and *θ* (the solid gray line) determines the cell size at division. Note that *c*_act_ and *θ* may have two intersections. Only the intersection on the left determines *V*_*d*_, because as the cell volume increases, *c*_act_ must rise to *θ*. There is no intersection beyond the critical threshold values (gray dashed lines). On the right is the cell size at birth *V*_*b*_ vs. *θ*. The dashed lines mark the predicted critical threshold values. (D) The cell integrates the information of one activator and one inhibitor using the AND logic. It divides once both *c*_act_ and *c*_inh_ reach their own thresholds. Here we show *V*_*b*_ vs. the activator’s threshold *θ*_act_ given a fixed threshold *θ*_inh_ = 350 *µ*m^-3^ for the inhibitor. In this figure, *K*_*n*,act_ = 12000 *µ*m^-3^, *K*_*n*,inh_ = 4000 *µ*m^-3^.

**S9 Fig. Imperfect sizer**. Simulations of *V*_*b*_ and *V*_*d*_. We simulate 5000 generations and trace one of the two daughter cells after division. For each generation, we first let the cell grow for a minimal time (20 min). We then use inverse transform sampling to generate the random variable *θ*, which is the activator-to-inhibitor ratio at cell division and has a probability density function *p*(*θ*) (see S1 Appendix for more details). In inverse transform sampling, we first take a random number *x* from a uniform distribution between 0 and 1. *θ* is then solved from *F* (*θ*) = *x*, where 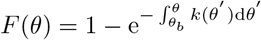 is the cumulative distribution function of *θ*. (A-B) *C*_1_ = 1, *C*_2_ = 2. The slope is similar to the imperfect sizer in [39, 42–44]. (C-D) *C*_1_ = 200, *C*_2_ = 1. The slope is similar to the near-adder in [45]. In this figure, *K*_*n*,act_ = 12000 *µ*m^-3^, *K*_*n*,inh_ = 4000 *µ*m^-3^.

**S10 Fig. The model of nondegradable inhibitor (multiple regulators)**. (A) Given a fixed *θ, V*_*b*_ increases after deleting one activator and decreases after deleting one inhibitor. The green and red dashed lines mark the range of *θ* that simultaneously allows WT, *activator* Δ and *inhibitor* Δ to divide. The black dashed line marks the linear relationship between *V*_*b*_ and *θ* in a wide range. *g*_act_ = *g*_inh_ = 10. (B) In the stochastic model, ⟨*V*_*b*_⟩ increases after deleting one activator and decreases after deleting one inhibitor. The CV of *V*_*b*_ changes mildly after deleting one regulator. (C) 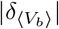 and |*δ*_CV_|vs. 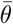 after deleting one activator. (D) 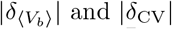 and |*δ*_CV_| vs. 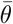 after deleting one inhibitor. In (B-D), the dashed lines mark the range of *θ* that simultaneously allows WT, *activator* Δ and *inhibitor* Δ to divide. (E) The distribution of cell size at birth *V*_*b*_ for WT and the mutant with the promoters of one activator and one inhibitor swapped. The average cell size and CV increase compared with the wild type, consistent with experiments [12]. 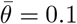. In this figure, *K*_*n*,act_ = 12000 *µ*m^-3^, *K*_*n*,inh_ = 1000 *µ*m^-3^, Δ*θ* = 0.009, *α* = 10.

**S11 Fig. Asymmetric division**. The cell divides asymmetrically with *γ* = 0.4. We track the daughter cell and simulate a single lineage to the steady state (see details in S1 Appendix). (A) Simulations and theoretical predictions of the cell size at birth *V*_*b*_ vs. the division threshold *θ* for the case of degradable regulators. (B) Simulations and theoretical predictions of *V*_*b*_ vs. *θ* for the case of degradable activator and nondegradable inhibitor. (C-G) The cell replicates its genes when the activator-to-inhibitor ratio rises to a threshold and then divides after a constant time *T*_*M*_ = 60 min. The inhibitor is nondegradable. (C) Simulations of *c*_act_ and *c*_inh_. The dashed lines mark cell division. (D) *c*_act_*/c*_inh_. (E) *V* . (F) *F*_*n*_ and *F*_*r*_. (G) *m*_inh_ and *p*_inh_. In (B-G), the inhibitor is distributed between the mother and daughter cells at division as Whi5 in budding yeast, with *η* = 0.45 [9]. In this figure, *K*_*n*,act_ = 12000 *µ*m^-3^, *K*_*n*,inh_ = 1000 *µ*m^-3^.

**S12 Fig. Simulations of multiple regulators with different weights and heterogeneous** *K*_*n,i*_ **(related to Fig 4)**. (A) In our model, the deletion of activators is equivalent to an increase in the division threshold *θ*. If the threshold change is small, e.g., *act1* Δ, the cell size at birth *V*_*b*_ is still proportional to the threshold (the yellow region) so that the CV of *V*_*b*_ remains constant. If the threshold change is significant, e.g., *act1* Δ *act2* Δ, the equivalent threshold is beyond the critical division threshold (the white region), and the cell becomes inviable (the red hollow circle). However, additional deletion of inh1 rescues the cell. (B) The distributions of *V*_*b*_ for WT, *act1* Δ and *act1* Δ *act2* Δ *inh1* Δ. (C) The deletion of inhibitors is equivalent to a decrease in the division threshold *θ*. If the threshold change is small, e.g., *inh3* Δ, *V*_*b*_ is still proportional to the threshold so that the CV of *V*_*b*_ remains constant. If the threshold change is significant, e.g., *inh2* Δ *inh3* Δ, *V*_*b*_ becomes a nonlinear function of the threshold (the purple region), and the CV increases. (D) The distributions of *V*_*b*_ for WT, *inh3* Δ, and *inh2* Δ *inh3* Δ. In (B, D), the mutants’ relative changes in average cell size and CV compared with WT are shown at the top. In simulations, we introduce different weights *χ*_act,*i*_ and *χ*_inh,*i*_ for different regulators. The cell divides once Σ_*i*_ *χ*_act,*i*_*c*_act,*i*_/Σ*τ*_*i*_*χ*_inh,*i*_*c*_inh,*i*_ = *θ* so that the change of average cell size can be different after deleting different activators or inhibitors. The *K*_*n*,act_ of 10 activators is an arithmetic sequence from 9000 to 15000 *µ*m^-3^. The *K*_*n*,inh_ of 10 inhibitors is an arithmetic sequence from 750 to 1250 *µ*m^-3^. Meanwhile, we also introduce heterogeneous *K*_*n,i*_ for other genes except the cell-cycle regulators, which follows a lognormal distribution with the mean *K*_*n*_ = 6000 *µ*m^-3^ and the CV equal to 0.5. Our conclusion regarding the CV of cell size after gene deletions remains valid in the presence of heterogeneous *K*_*n,i*_. *χ*_act,1_ = *χ*_act,2_ = 2, *χ*_act,3_ = … = *χ*_act,6_ = 1, *χ*_act,7_ = … = *χ*_act,10_ = 0.5. *χ*_inh,1_ = *χ*_inh,2_ = 2, *χ*_inh;3_ = … = *χ*_nh;6_ = 1, *χ*_inh;7_ = … = *χ*_inh;10_ = 0:5. 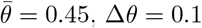, Δ*θ*= 0.1.

## Acknowledgments

We thank Lingyu Meng, Qirun Wang, Yiyang Ye, and Yichen Yan for helpful discussions related to this work, and three anonymous reviewers for constructive comments. The research was funded by National Key R&D Program of China (2021YFF1200500) and supported by grants from Peking-Tsinghua Center for Life Sciences.

