## Supplementary Material for "Implications of differential size-scaling of cell-cycle regulators on cell size homeostasis"

June 2, 2023

**1** Yuanpei College, Peking University, Beijing, China

**2** Center for Quantitative Biology, Academy for Advanced Interdisciplinary Studies, Peking University, Beijing, China

**3** Peking-Tsinghua Center for Life Sciences, Academy for Advanced Interdisciplinary Studies, Peking University, Beijing, China

*

#### 1 Detailed analysis of the drop in $c_{\mathrm{act}}/c_{\mathrm{inh}}$ right after division

Because the cell-cycle regulators are degradable, their numbers change much faster than $F_n$ and $F_r$. Therefore, $F_n(t) \approx F_n(0)$ and $F_r(t) \approx F_r(0)$ right after cell division. For the activator ($i = 3$) and the inhibitor ($i = 4$),

$$\frac{\mathrm{d}m_i}{\mathrm{d}t} = \Gamma_n g_i \frac{F_n(0)c_n}{F_n(0)c_n + K_{n,i}} - \frac{m_i}{\tau_m}, \tag{S1}$$

$$\frac{\mathrm{d}p_i}{\mathrm{d}t} = \Gamma_r m_i \frac{F_r(0)c_r}{F_r(0)c_r + K_r} - \frac{p_i}{\tau_p}. \tag{S2}$$

The solution is

$$p_i(t) = p_i(0)\mathrm{e}^{-\frac{t}{\tau_p}} + \frac{B\tau_m\tau_p}{\tau_p - \tau_m}\left(m_i(0)(\mathrm{e}^{-\frac{t}{\tau_p}} - \mathrm{e}^{-\frac{t}{\tau_m}}) + A_i(\tau_m \mathrm{e}^{-\frac{t}{\tau_m}} - \tau_p \mathrm{e}^{-\frac{t}{\tau_p}} + \tau_p - \tau_m)\right), \tag{S3}$$

where

$$A_i = \Gamma_n g_i \frac{F_n(0)c_n}{F_n(0)c_n + K_{n,i}}, \quad B = \Gamma_r \frac{F_r(0)c_r}{F_r(0)c_r + K_r}. \tag{S4}$$

From Eq S3 and $K_{n,\mathrm{act}} > K_{n,\mathrm{inh}}$, we get

$$\frac{\mathrm{d}}{\mathrm{d}t}\left(\frac{p_{\mathrm{act}}}{p_{\mathrm{inh}}}\right) = (A_{\mathrm{act}} - A_{\mathrm{inh}})C(t) < 0, \tag{S5}$$

where

$$C(t) = \frac{B\tau_m\left(p_i(0)(\tau_m - \tau_p)(\mathrm{e}^{\frac{t}{\tau_m}} - 1) + B\tau_p m_i(0)(\tau_p(\mathrm{e}^{\frac{t}{\tau_p}} - 1) - \tau_m(\mathrm{e}^{\frac{t}{\tau_m}} - 1))\right)}{(\tau_m - \tau_p)\mathrm{e}^{\frac{t}{\tau_m} + \frac{t}{\tau_p}} p_{\mathrm{inh}}^2} > 0. \tag{S6}$$

Therefore, $c_{\mathrm{act}}/c_{\mathrm{inh}}$ decreases right after cell birth. Additionally, from Eq S3, we find that in the limit of very small $\tau_m$ and $\tau_p$, the decrease of $c_{\mathrm{act}}/c_{\mathrm{inh}}$ is very sharp. A similar analysis applies to the modified model with gene replication in Section 4 of S1 Appendix.

#### 2 The stability analysis of the cell cycle

In this section, we prove the stability of the cell cycle in the deterministic model. When the ratio of the activator to the inhibitor reaches the threshold value, the system is on the $2N-1$ dimensional hyperplane $\theta = \frac{p_{\mathrm{act}}}{p_{\mathrm{inh}}}$ in the phase space, where $N$ is the total number of genes. Right after division, it is still on this hyperplane because all mRNAs and proteins halve. Without loss of generality, we use the quantities right after division to describe the system's state. We denote $\boldsymbol{x}_k = (m_{1,k}, ..., m_{N,k}, n_k, p_{\mathrm{inh},k}, r_k, p_{5,k}, ..., p_{N,k})^T$ as the state vector of the system when it reaches this hyperplane for the $k$-th time. Note that we exclude $p_{\mathrm{act},k}$ from the state vector for it is identically equal to $\theta p_{\mathrm{inh},k}$ on the hyperplane. The Poincaré map [1] $P$ satisfies $\boldsymbol{x}_k = P(\boldsymbol{x}_{k-1})$. In the following we prove that the Jacobian matrix of $P$ only has eigenvalues with absolute values smaller than 1.

For nondegradable proteins, the solution to Eq 8 is

$$p_i(t) = p_i(0) + \Gamma_{n,i} g_i \tau_m \Gamma_r \int_0^t \frac{F_n(s)c_n}{F_n(s)c_n + K_n} \frac{F_r(s)c_r}{F_r(s)c_r + K_r} \mathrm{d}s, \tag{S7}$$

In particular,

$$n(t) = n(0) + \Gamma_{n,n} g_n \tau_m \Gamma_r \int_0^t \frac{F_n(s)c_n}{F_n(s)c_n + K_n} \frac{F_r(s)c_r}{F_r(s)c_r + K_r} \mathrm{d}s, \tag{S8}$$

Therefore,

$$p_i(t) = p_i(0) + \frac{\Gamma_{n,i} g_i}{\Gamma_{n,n} g_n}(n(t) - n(0)). \tag{S9}$$

According to Eqs 20,21,

$$m_{i,k} = \Gamma_{n,i} g_i \tau_m \tfrac{\tilde{K}}{2((1-\theta)K_n + \tilde{K})}, \quad k \in \mathbb{N}_+, \tag{S10}$$

$$n_k = \tfrac{n_c c_n (1-\theta)\tilde{K}}{2((1-\theta)c_n - \tilde{K})((1-\theta)K_n + \tilde{K})}, \quad k \in \mathbb{N}_+. \tag{S11}$$

For proteins except RNAPs and cell-cycle regulators, from Eq S9,

$$2p_{i,k} = p_{i,k-1} + \frac{\Gamma_{n,i} g_i}{\Gamma_{n,n} g_n}(2n_k - n_{k-1}) = p_{i,k-1} + \frac{\Gamma_{n,i} g_i}{\Gamma_{n,n} g_n} \frac{n_c c_n (1-\theta)\tilde{K}}{2((1-\theta)c_n - \tilde{K})((1-\theta)K_n + \tilde{K})}, \quad k \in \mathbb{N}_+. \tag{S12}$$

From Eq 16,

$$2p_{\mathrm{inh},k} = \Gamma_n g_{\mathrm{inh}} \tau_m \Gamma_r \tau_p \frac{F_{n,d} c_n}{F_{n,d} c_n + K_{n,\mathrm{inh}}} \frac{F_{r,d} c_r}{F_{r,d} c_r + K_r}, \tag{S13}$$

where the fraction of free RNAPs at cell division $F_{n,d}$ is a constant (Eq 17), while the fraction of free ribosomes at cell division $F_{r,d}$ depends on $r_{k-1}$ (Eq 9 and Eq S12 about $r_k$). Therefore, from Eqs S10-S13, $\frac{\partial \boldsymbol{x}_k}{\partial \boldsymbol{x}_{k-1}}$ is a upper-triangular matrix with diagonal entries $\underbrace{0, ..., 0}_{N+2}, \underbrace{\frac{1}{2}, ..., \frac{1}{2}}_{N-3}$. The eigenvalues of the Jacobian matrix are precisely these diagonal entries, and their absolute values are all small than 1. Therefore, the periodic orbit is stable against infinitesimal perturbation.

#### 3 Imperfect sizer

To accommodate the fact that cells display imperfect sizers [2, 3, 4, 5, 6, 7, 8], we assume that cell division is a stochastic process with its probability increases with the activator-to-inhibitor ratio. This idea is similar to some previous work [9, 10, 11, 3, 12]. Specifically, we first let the cell grow for a short period in order to set the minimal cell-cycle time and avoid the transient effect right after division. After this period, the probability for a cell with the activator-to-inhibitor ratio in the infinite interval $[\theta, \theta + \mathrm{d}\theta)$ to divide is $k(\theta)\mathrm{d}\theta$. The probability distribution for the ratio $\theta$ at division is

$$p(\theta) = k(\theta)\mathrm{e}^{-\int_{\theta_b}^{\theta} k(\theta^{'})\mathrm{d}\theta^{'}}, \tag{S14}$$

where $\theta_b$ is the activator-to-inhibitor ratio right after the short period. To ensure $\int_{\theta_b}^{\theta_2} p(\theta)\mathrm{d}\theta = 1$, where $\theta_2$ is the larger critical threshold value in the main text, $k(\theta)$ must satisfy $\lim_{\theta \to \theta_2} k(\theta) = +\infty$.

The cell volume $V$ can be expressed by an increasing function $V(\theta)$, because the ratio $\theta$ between the regulators undergoing size-dependent expression is a measurement for cell volume. The expectation for the division size $V_d$ is then

$$\langle V_d \rangle = \int_{\theta_b}^{\theta_2} V(\theta)p(\theta)\mathrm{d}\theta = \int_{\theta_b}^{\theta_2} V(\theta)k(\theta)\mathrm{e}^{-\int_{\theta_b}^{\theta} k(\theta^{'})\mathrm{d}\theta^{'}}\mathrm{d}\theta. \tag{S15}$$

We consider a cell born with a larger size. The expectation for its division size $V_d^{'}$ is then

$$\langle V_d^{'} \rangle = \int_{\theta_b^{'}}^{\theta_2} V(\theta)k(\theta)\mathrm{e}^{-\int_{\theta_b^{'}}^{\theta} k(\theta^{'})\mathrm{d}\theta^{'}}\mathrm{d}\theta, \tag{S16}$$

where $\theta_b^{'} > \theta_b$. If $k(\theta)$ becomes nonzero before $\theta$ reaches $\theta_b^{'}$,

$$\begin{aligned}
\langle V_d^{'} \rangle - \langle V_d \rangle &= \int_{\theta_b^{'}}^{\theta_2} V(\theta)k(\theta)\left(\mathrm{e}^{-\int_{\theta_b^{'}}^{\theta} k(\theta^{'})\mathrm{d}\theta^{'}} - \mathrm{e}^{-\int_{\theta_b}^{\theta} k(\theta^{'})\mathrm{d}\theta^{'}}\right)\mathrm{d}\theta - \int_{\theta_b}^{\theta_b^{'}} V(\theta)k(\theta)\mathrm{e}^{-\int_{\theta_b}^{\theta} k(\theta^{'})\mathrm{d}\theta^{'}}\mathrm{d}\theta \\
&> V(\theta_b^{'})\int_{\theta_b^{'}}^{\theta_2} k(\theta)\left(\mathrm{e}^{-\int_{\theta_b^{'}}^{\theta} k(\theta^{'})\mathrm{d}\theta^{'}} - \mathrm{e}^{-\int_{\theta_b}^{\theta} k(\theta^{'})\mathrm{d}\theta^{'}}\right)\mathrm{d}\theta - V(\theta_b^{'})\int_{\theta_b}^{\theta_b^{'}} k(\theta)\mathrm{e}^{-\int_{\theta_b}^{\theta} k(\theta^{'})\mathrm{d}\theta^{'}}\mathrm{d}\theta \\
&= V(\theta_b^{'})\left(1 - \int_{\theta_b^{'}}^{\theta_2} k(\theta)\mathrm{e}^{-\int_{\theta_b}^{\theta} k(\theta^{'})\mathrm{d}\theta^{'}}\mathrm{d}\theta\right) - V(\theta_b^{'})\int_{\theta_b}^{\theta_b^{'}} k(\theta)\mathrm{e}^{-\int_{\theta_b}^{\theta} k(\theta^{'})\mathrm{d}\theta^{'}}\mathrm{d}\theta = 0,
\end{aligned} \tag{S17}$$

which means $\langle V_d \rangle$ increases with $\theta_b$. Therefore, $\langle V_d \rangle$ increases with the birth volume, which is the imperfect sizer. Intuitively, the cell with a smaller birth size is more likely to divide at a smaller size because it has a finite probability to divide before it reaches the birth size of the larger cell.

As a simple example, we assume

$$k(\theta) = \begin{cases} C_1\left(\frac{\theta-\theta_1}{\theta_2-\theta}\right)^{C_2}, & \theta_1 < \theta < \theta_2 \\ 0, & \text{others} \end{cases}. \tag{S18}$$

where the constant $C_1, C_2 > 0$ and $\theta_1$ is the smaller critical threshold value in the main text. We numerically generate imperfect sizers with the slopes similar to experiments (S9 Fig). Our results show that, in principle, differential scaling mechanisms can enact a broad spectrum of cell size control strategies. We would like to clarify that the phenomenological adder (or near-adder) over the entire cell cycle observed in eukaryotic cells [13, 7] can emerge from the combination of independent regulations at different cell-cycle stages [3, 14, 15].

Finally, we want to mention that for any distribution $p(\theta)$, there exists a corresponding distribution $k(\theta)$. Integrating over both sides of Eq S14, we get

$$\int_{\theta_b}^{x} p(\theta)\mathrm{d}\theta = 1 - \mathrm{e}^{-\int_{\theta_b}^{x} k(\theta^{'})\mathrm{d}\theta^{'}}, \tag{S19}$$

$$\int_{\theta_b}^{x} k(\theta^{'})\mathrm{d}\theta^{'} = -\ln\left(1 - \int_{\theta_b}^{x} p(\theta)\mathrm{d}\theta\right). \tag{S20}$$

Taking the derivative on both sides of Eq S20, we can determine $k(\theta)$ from $p(\theta)$,

$$k(\theta) = \frac{p(\theta)}{1 - \int_{\theta_b}^{\theta} p(\theta')\mathrm{d}\theta'}. \tag{S21}$$

### 4 The modified model with gene replication

In this section, we assume that when the ratio of the activator to the inhibitor $c_{\mathrm{act}}/c_{\mathrm{inh}}$ rises to the threshold $\theta$, the cell replicates its genes instantly and divides after a constant time $T_M$. Like the simplified model in the main text, here the cell can also reach a periodic steady state (S6A-C Fig). The difference is that here when genes replicate, the free RNAP fraction $F_n$ drops abruptly (S6D Fig), causing the reduction of $c_{\mathrm{act}}/c_{\mathrm{inh}}$.

We can easily obtain the expression of $V_b$ after some modification to the corresponding deviation in Methods. We use subscript $S$ to denote the values at gene replication. Substituting Eqs 15-16 into

$$\theta = \frac{p_{\mathrm{act},S}}{p_{\mathrm{inh},S}}, \tag{S22}$$

we obtain

$$F_{n,S} = \frac{\theta K_{n,\mathrm{act}} - K_{n,\mathrm{inh}}}{(1-\theta)c_n}. \tag{S23}$$

Substituting Eq S23 into Eq 18, we get the critical size [16] to enter S phase

$$V_S = \tfrac{a}{c_n} n_S = \tfrac{a n_c (1-\theta)\tilde{K}}{((1-\theta)c_n - \tilde{K})((1-\theta)K_n + \tilde{K})}, \tag{S24}$$

$$V_b = \tfrac{1}{2} V_S \mathrm{e}^{\mu T_M} = \tfrac{\mathrm{e}^{\mu T_M} a n_c (1-\theta)\tilde{K}}{2((1-\theta)c_n - \tilde{K})((1-\theta)K_n + \tilde{K})}. \tag{S25}$$

When $T_M = 0$, Eq S25 becomes the same as Eq 2 in the main text. Noticing the constrain of $0 < F_{n,S} < 1$ in Eq S23, we get the range for $\theta$, which is the same as Eq 3.

We estimate the growth rate $\mu$ in Eq S25 as follows. Substituting Eq 13 about $r$ and Eq 18 into Eq 33, we get

$$\frac{F_r c_r}{F_r c_r + K_r} \approx \frac{n_c \Gamma_{n,r} g_r (1 - F_r)}{\Gamma_{n,n} g_n \tau_m (1 - F_n) \sum_i \Gamma_{n,i} g_i \left(1 + \frac{\Gamma_r L_i}{v_r}\right)}. \tag{S26}$$

When $F_n \ll 1$, Eq S26 becomes

$$\frac{F_r c_r}{F_r c_r + K_r} \approx \frac{n_c \Gamma_{n,r} g_r (1 - F_r)}{\Gamma_{n,n} g_n \tau_m \sum_i \Gamma_{n,i} g_i \left(1 + \frac{\Gamma_r L_i}{v_r}\right)}, \tag{S27}$$

from which we can determine $F_r$. Then, from Eq 34, we obtain the growth rate $\mu$. The constant $\mu$ means that the cell grows exponentially. This scenario is called Phase 1 of gene expression [17]. Although a large cell deviates from Phase 1, its growth rate changes relatively mildly, so the $\mu$ calculated above is still a good approximation. Comparison between the predictions with simulations shows good agreement (S6E Fig).

### 5 The case of degradable activator and nondegradable inhibitor

Although there are many short-lived cell-cycle regulators, such as the activator Cln3 [18, 19] and the inhibitor Sic1 [20, 21, 22] in budding yeast, the inhibitor Whi5 degrades slowly [23, 24]. In mammalian cells, the cell-cycle inhibitor Rb is also very stable [25, 26, 27]. Therefore, the case of degradable activator but nondegradable inhibitor is also worth discussion. We set the transcription initiation rate of the inhibitor to be $\Gamma_n/\alpha$ ($\alpha > 1$) so that the level of activators and inhibitors are more comparable.

#### 5.1 Derivation of the cell size at birth and the two critical threshold values

Combining Eqs 8,11, we get

$$\frac{\mathrm{d}}{\mathrm{d}t}\left(\frac{p_{\mathrm{inh}}}{n}\right) = \frac{\Gamma_{n,n} g_n \tau_m \Gamma_r}{n} \frac{F_n c_n}{F_n c_n + K_n} \frac{F_r c_r}{F_r c_r + K_r} \left(\frac{\Gamma_n g_{\mathrm{inh}}}{\alpha \Gamma_{n,n} g_n} \frac{F_n c_n + K_n}{F_n c_n + K_{n,\mathrm{inh}}} - \frac{p_{\mathrm{inh}}}{n}\right). \tag{S28}$$

We use quasi-steady-state approximation for $\frac{p_{\mathrm{inh}}}{n}$ at division: $\frac{\mathrm{d}}{\mathrm{d}t}\left(\frac{p_{\mathrm{inh}}}{n}\right) = 0$. Therefore,

$$p_{\mathrm{inh},d} = \frac{\Gamma_n g_{\mathrm{inh}}}{\alpha \Gamma_{n,n} g_n} \frac{F_{n,d} c_n + K_n}{F_{n,d} c_n + K_{n,\mathrm{inh}}} n_d. \tag{S29}$$

In the simple case of $g_{\mathrm{act}} = g_{\mathrm{inh}} = 1$, substituting Eqs 15,S29 into Eq 1, and using Eq 18, we get

$$\frac{\theta}{\alpha} = \frac{\Gamma_{n,n} g_n \tau_m \Gamma_r \tau_p}{n_c} \frac{(1 - F_{n,d})(F_{n,d} c_n + K_{n,\mathrm{inh}})}{F_{n,d} c_n + K_{n,\mathrm{act}}} \frac{F_{r,d} c_r}{F_{r,d} c_r + K_r}. \tag{S30}$$

From Eqs S26,S30, we can numerically solve $F_{n,d}$, and eventually get $n_b$ and $V_b$ (S7A Fig, Prediction 1).

Experimentally, cell growth in an unperturbed cell cycle is typically nearly exponential [28, 29, 30], so we are especially concerned about the cell in Phase 1. According to [31], for the cell in Phase 1,

$$p_{\mathrm{act}}(t) = \frac{\Gamma_n K_n n_c}{\Gamma_{n,n}} \frac{\mu \tau_p n(0) \mathrm{e}^{\mu t}}{K_{n,\mathrm{act}} n_c - (K_{n,\mathrm{act}} - K_n) n(0) \mathrm{e}^{\mu t}}, \tag{S31}$$

$$p_{\mathrm{inh}}(t) = p_{\mathrm{inh}}(0) + \frac{\Gamma_n K_n n_c}{\alpha \Gamma_{n,n}} \frac{1}{K_n - K_{n,\mathrm{inh}}} \ln \frac{K_{n,\mathrm{inh}} n_c - (K_{n,\mathrm{inh}} - K_n) n(0) \mathrm{e}^{\mu t}}{K_{n,\mathrm{inh}} n_c - (K_{n,\mathrm{inh}} - K_n) n(0)}, \tag{S32}$$

where $\mu$ is estimated as Section 4 of S1 Appendix. Using $p_{\mathrm{inh},d} = 2 p_{\mathrm{inh},b}$, we get

$$p_{\mathrm{inh},d} = \frac{2 \Gamma_n K_n n_c}{\alpha \Gamma_{n,n}} \frac{1}{K_n - K_{n,\mathrm{inh}}} \ln \frac{K_{n,\mathrm{inh}} n_c - 2(K_{n,\mathrm{inh}} - K_n) n_b}{K_{n,\mathrm{inh}} n_c - (K_{n,\mathrm{inh}} - K_n) n_b}, \tag{S33}$$

Substituting Eqs S31,S33 into Eq 1, we obtain

$$\frac{\theta}{\alpha} = \frac{\frac{\mu \tau_p n_b}{K_{n,\mathrm{act}} n_c - 2(K_{n,\mathrm{act}} - K_n) n_b}}{\frac{1}{K_n - K_{n,\mathrm{inh}}} \ln \frac{K_{n,\mathrm{inh}} n_c - 2(K_{n,\mathrm{inh}} - K_n) n_b}{K_{n,\mathrm{inh}} n_c - (K_{n,\mathrm{inh}} - K_n) n_b}}. \tag{S34}$$

Eq S34 can implicitly determine $n_b$ and $V_b$. To get an approximate explicit solution for $n_b$, we use Taylor expansion

$$\begin{aligned} \frac{1}{K_n - K_{n,\mathrm{inh}}} \ln \frac{K_{n,\mathrm{inh}} n_c - 2(K_{n,\mathrm{inh}} - K_n) n_b}{K_{n,\mathrm{inh}} n_c - (K_{n,\mathrm{inh}} - K_n) n_b} &= \frac{1}{K_n - K_{n,\mathrm{inh}}} \ln \left(1 + \frac{(K_n - K_{n,\mathrm{inh}}) n_b}{K_{n,\mathrm{inh}} n_c - (K_{n,\mathrm{inh}} - K_n) n_b}\right) \\ &\approx \frac{n_b}{K_{n,\mathrm{inh}} n_c - (K_{n,\mathrm{inh}} - K_n) n_b}, \end{aligned} \tag{S35}$$

so we get

$$n_b \approx \frac{n_c (\theta K_{n,\mathrm{act}}/\alpha - \mu \tau_p K_{n,\mathrm{inh}})}{2\theta (K_{n,\mathrm{act}} - K_n)/\alpha + \mu \tau_p (K_n - K_{n,\mathrm{inh}})}, \tag{S36}$$

Finally, we obtain $V_b$ (S7A Fig, Prediction 2), which fits well with simulations (S7A Fig) for relatively small $V_b$. The deviation for large $V_b$ is reasonable, because Eqs S31-S32 are no longer accurate after the cell leaves Phase 1. Letting $n_b \to 0$ in Eq S34 or Eq S36, we find that one of the critical thresholds is $\theta_1 = \frac{\alpha\mu\tau_p K_{n,\mathrm{inh}}}{K_{n,\mathrm{act}}}$. It's hard to analytically calculate the other critical point $\theta_2$, but S7B Fig provides an illustration. When the cell is excessively large, genes and mRNAs get saturated, so the number of the degradable activator becomes constant, and the number of the nondegradable inhibitor is proportional to cell volume. Therefore, $c_{\mathrm{act}}/c_{\mathrm{inh}}$ approaches to zero, and the cell cannot divide any more. The phenomenon that large cell volume prevents cell division is consistent with experiments [32, 33, 34].

##### 5.2 The correlations between $V_b$ and $V_d$

As in the main text, we consider the noise in the threshold value. Given a $\theta$, from Eqs 18,S26,S30 we can determine the number of RNAPs at cell division $n_d$ and then the cell size at division $V_d$ approximately. Therefore, $V_d$ in a certain generation nearly only depends on the threshold of that generation and is consequently uncorrelated with $V_b$. This prediction is close to simulations (S7C-E Fig) and experiments [2, 8].

#### 6 Alternative criteria for cell division

In this section, we explore alternative criteria that govern cell-cycle progression. We first assume that the cell divides once the concentration $c_i$ of one activator or inhibitor $i$ reaches a threshold value $\theta_i$. When this event happens,

$$c_i = \frac{p_i}{V} = \frac{c_n p_i}{an} = \theta_i. \tag{S37}$$

The cell volume at birth $V_{b,i} = c_i^{-1}(\theta_i)/2$, where $c_i^{-1}$ denotes the inverse function of $c_i(V)$ (S8 Fig).

If the regulator $i$ is degradable, substituting Eq 15 or Eq 16 into Eq S37, and using Eq 18, we get

$$\tilde{\theta}_i = \frac{\Gamma_{n,n} g_n \tau_m \Gamma_r \tau_p}{n_c} \frac{(1-F_{n,d})(F_{n,d}c_n + K_n)}{F_{n,d}c_n + K_{n,i}} \frac{F_{r,d}c_r}{F_{r,d}c_r + K_r}. \tag{S38}$$

where $\tilde{\theta}_i \equiv \frac{a\Gamma_{n,n}g_n}{c_n\Gamma_{n,i}g_i}\theta_i$. From Eqs S26,S38, we can numerically solve $F_{n,d}$, and eventually get $V_b$. The derivation here is similar to Section 5 of S1 Appendix, and we can also obtain the smaller critical threshold $\tilde{\theta}_{i,1} = \frac{\mu\tau_p K_n}{K_{n,i}}$ (S8A-B Fig).

If the regulator $i$ is nondegradable, substituting Eq S29 into Eq S37, we get

$$F_{n,d} = \frac{\tilde{\theta}_i K_{n,i} - K_n}{(1-\tilde{\theta}_i)c_n}. \tag{S39}$$

Similar to the derivation in Methods, we obtain the cell volume at birth (S8C Fig)

$$V_{b,i} = \frac{an_c(1-\tilde{\theta}_i)(\tilde{\theta}_i K_{n,i} - K_n)}{2((1-\tilde{\theta}_i)c_n - (\tilde{\theta}_i K_{n,i} - K_n))((1-\tilde{\theta}_i)K_n + (\tilde{\theta}_i K_{n,i} - K_n))}, \tag{S40}$$

where

$$\frac{K_n}{K_{n,i}} < \tilde{\theta}_i < \frac{c_n + K_n}{c_n + K_{n,i}}. \tag{S41}$$

We then consider the logic gates for multiple regulators motivated by the AND gate logic observed in *E. coli* [35, 36]. For the AND gate, i.e., the cell divides after each $c_i$ reaches its own threshold $\theta_i$, the birth volume $V_{b,\mathrm{AND}} = \max\{V_{b,i}\}$ (S8D Fig). For the OR gate, i.e., the cell divides once one of the $c_i$ reaches its threshold, the birth volume $V_{b,\mathrm{OR}} = \min\{V_{b,i}\}$. Similar methods can be applied to the combination of AND and OR gates. We note that the combination of logic gates cannot explain the phenotype of some mutants. For example, *cln3*$\Delta$*bck2*$\Delta$ for budding yeast is lethal, while *cln3*$\Delta$*bck2*$\Delta$*whi5*$\Delta$ is viable [37], which we think is beyond a combination of logic gates.

### 7 Asymmetric division

In this section, we consider asymmetric division to better apply our model to asymmetrically dividing organisms such as budding yeast [38]. The proportion of the cell volume the daughter cell inherits is $\gamma < 1/2$.

For the case of degradable activator and inhibitor, using $n_b = \gamma n_d$ and Eq 21, we easily get the cell volume at birth for the daughter cell

$$V_b = \frac{\gamma a n_c (1-\theta)\tilde{K}}{((1-\theta)c_n - \tilde{K})((1-\theta)K_n + \tilde{K})}. \tag{S42}$$

The two critical thresholds remain the same (S11A Fig).

We also consider a modified model in which the inhibitor is nondegradable and some inhibitor molecules are bound to chromatins at cell division as Whi5 in budding yeast [23, 24, 39]. Therefore, the proportion $\eta$ of the inhibitor molecules that the daughter cell inherits satisfies $\gamma < \eta < 1/2$. Similar to derivations in Section 5 of S1 Appendix, using $n_b = \gamma n_d$ and $p_{\mathrm{inh},b} = \eta p_{\mathrm{inh},d}$ instead, we obtain

$$n_b \approx \frac{n_c\left(\frac{(1-\gamma)\theta}{(1-\eta)\alpha}K_{n,\mathrm{act}} - \mu\tau_p K_{n,\mathrm{inh}}\right)}{\frac{1}{\gamma}\frac{(1-\gamma)\theta}{(1-\eta)\alpha}(K_{n,\mathrm{act}} - K_n) + \mu\tau_p(K_n - K_{n,\mathrm{inh}})}. \tag{S43}$$

When $\gamma = \eta = 1/2$, Eq S43 becomes Eq S36. The smaller critical threshold $\theta_1 = \frac{(1-\eta)\alpha\mu\tau_p K_{n,\mathrm{inh}}}{(1-\gamma)K_{n,\mathrm{act}}}$. Simulations confirm our predictions (S11B Fig).

In the following, we assume that the cell replicates its genes when the activator-to-inhibitor ratio rises to a threshold and then divides after a constant time $T_M$. The cell can reach a periodic steady state (S11C-E Fig). The difference is that here both when genes replicate and when the cell divides, the free RNAP fraction $F_n$ drops abruptly (S11F Fig), causing the reduction of $c_{\mathrm{act}}/c_{\mathrm{inh}}$. Due to the chromatin-based partitioning of the inhibitor, $c_{\mathrm{inh}}$ increases abruptly at cell birth (S11C Fig). Interestingly, the mRNA number and protein synthesis rate of the inhibitor increase after G1/S transition (S11G Fig), which is similar to the behavior of Whi5 synthesis in budding yeast [23, 24]. We remark that our model predicts that the *WHI5* mRNA number and Whi5 synthesis rate in S/G2/M is approximately twice as much as that in G1 because of the doubled *WHI5* gene copy number and the strong affinity of the *WHI5* promoter with RNAP. This prediction falls between the two opposite views ( [40] vs. [41, 42]). Specific regulation on Whi5 synthesis seems to exist in different cell-cycle stages. We acknowledge that we cannot quantitatively compare the oscillation amplitude in our model with that of Whi5, given that the behavior and regulation of Whi5 synthesis remains an open question.

In summary, asymmetric division does not lead to qualitative difference in our results, and our methodology can be applied to this case with minor modifications.

Table S1: A summary of the parameters used in the simulations if not mentioned in the text. Note that some of the parameters may not be realistic estimations of any specific organisms and our main conclusions are independent of the chosen parameters.

| Variables | Meaning | Values |
|---|---|---|
| $K_n$ | average transcriptional MM constant | 6000 $\mu$m$^{-3}$ |
| $K_r$ | translational MM constant | 6000 $\mu$m$^{-3}$ |
| $v_n$ | RNAP elongation speed | 36 nt/s |
| $v_r$ | ribosome elongation speed | 12 aa/s |
| $g_n$ | copy number of the RNAP gene | 1 |
| $g_r$ | copy number of the ribosome gene | 5 |
| $g_i$ $(i > 2)$ | copy number of other genes | 1 |
| $L_n$ | number of amino acids of the RNAP | $10^3$ aa |
| $L_r$ | number of amino acids of the ribosome | $10^4$ aa |
| $L$ | number of amino acids of other proteins | 500 aa |
| $\rho$ | protein mass per cell volume | $10^{10}$ aa/$\mu$m$^{-3}$ |
| $a$ | ratio between cell volume and nuclear volume | 20 |
| $N$ | total number of genes | 2000 |
| $\Gamma_r$ | translational initiation rate | 10 min$^{-1}$ |
| $\tau_m$ | lifetime of mRNAs | 5 min |
| $\tau_p$ | lifetime of degradable proteins | 5 min |
| - | lifetime of nondegradable proteins | $10^4$ min |
| $T_M$ | time between gene replication and cell division | 60 min |

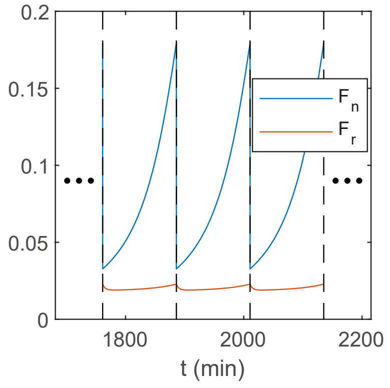


S1 Fig.

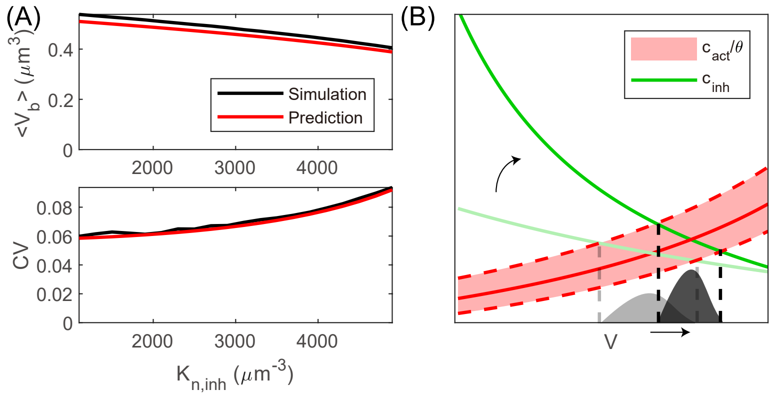


S2 Fig.

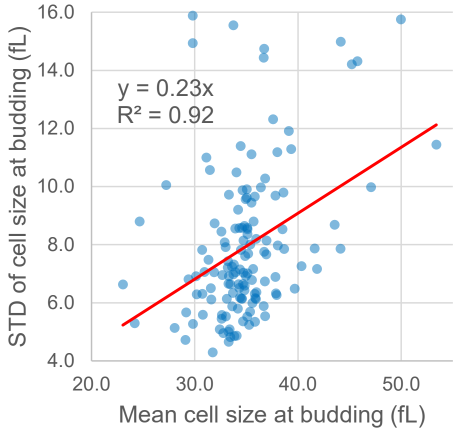


S3 Fig.

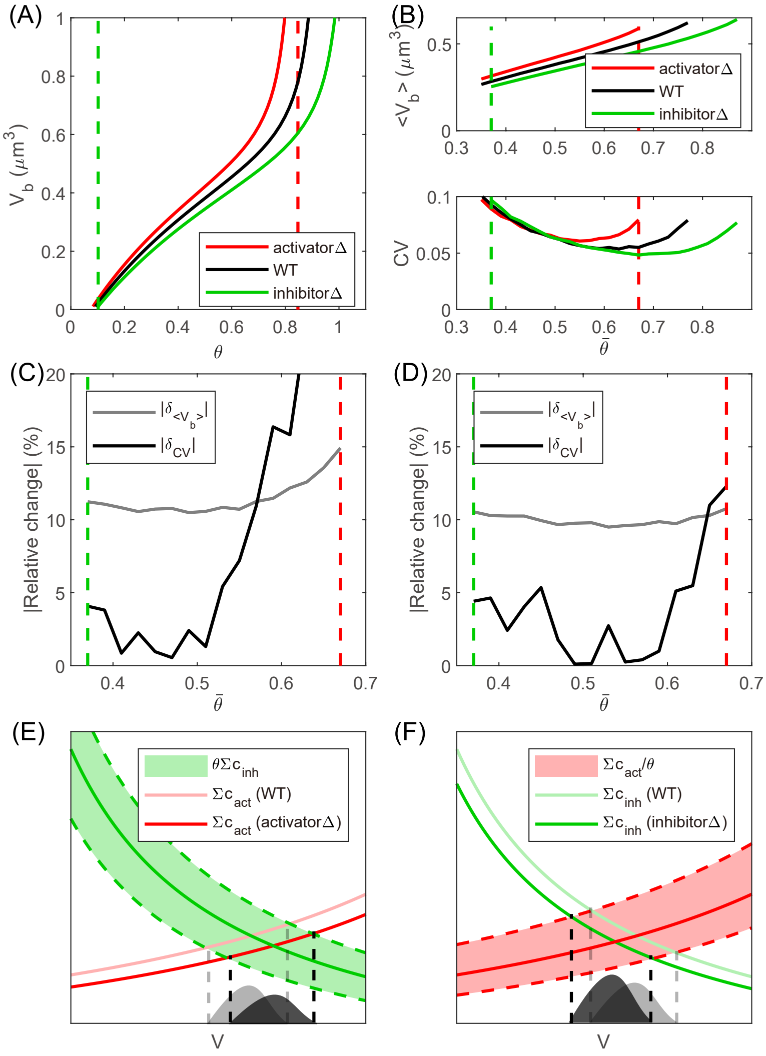

(A)
(B)
(C)
(D)
(E)
(F)
activatorΔ
WT
inhibitorΔ
|Relative change| (%)
|δ_CV|
θΣc_inh
Σc_act (WT)
Σc_act (activatorΔ)
Σc_act/θ
Σc_inh (WT)
Σc_inh (inhibitorΔ)
V


S4 Fig.

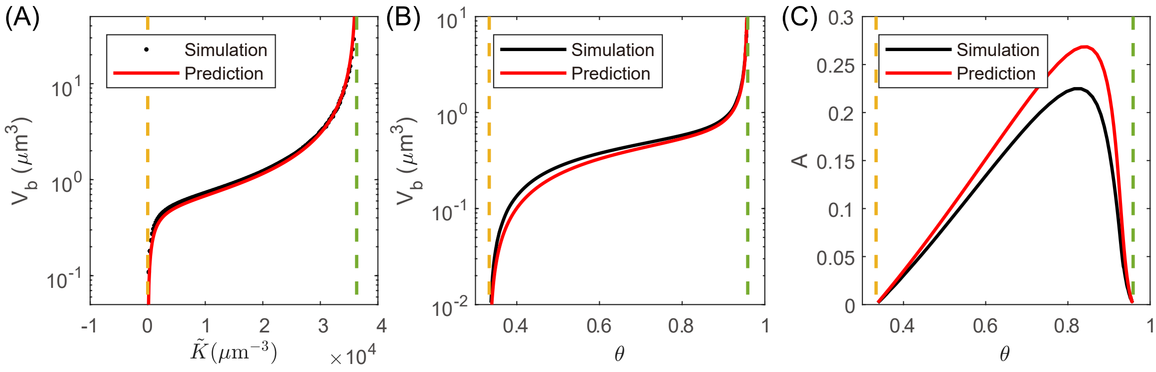

(A)
(B)
(C)
Simulation
Prediction


S5 Fig.

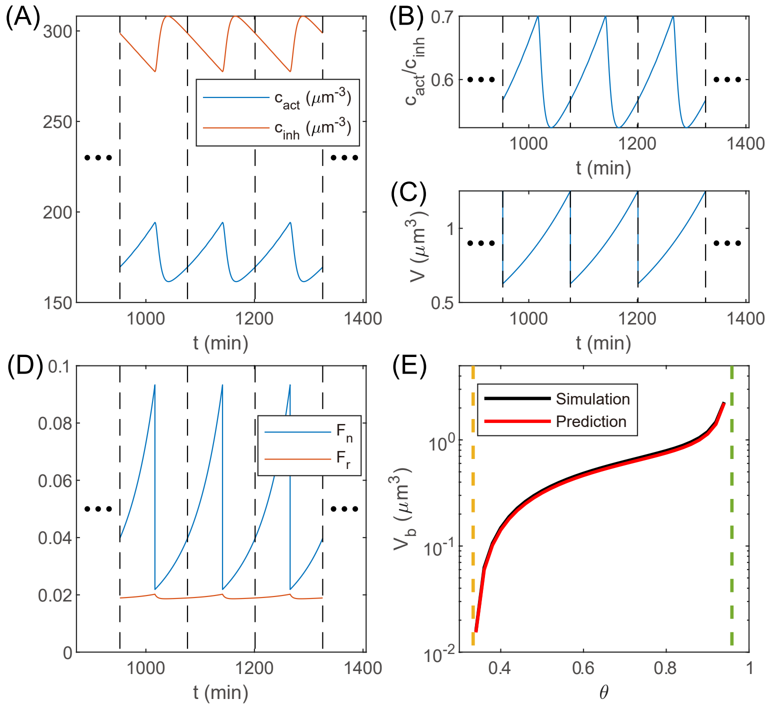
(A)
(B)
(C)
(D)
(E)
$c_{act}$ ($\mu$m$^{-3}$)
$c_{inh}$ ($\mu$m$^{-3}$)
t (min)
$c_{act}/c_{inh}$
V ($\mu$m$^3$)
$F_n$
$F_r$
Simulation
Prediction
$V_b$ ($\mu$m$^3$)
$\theta$

S6 Fig.

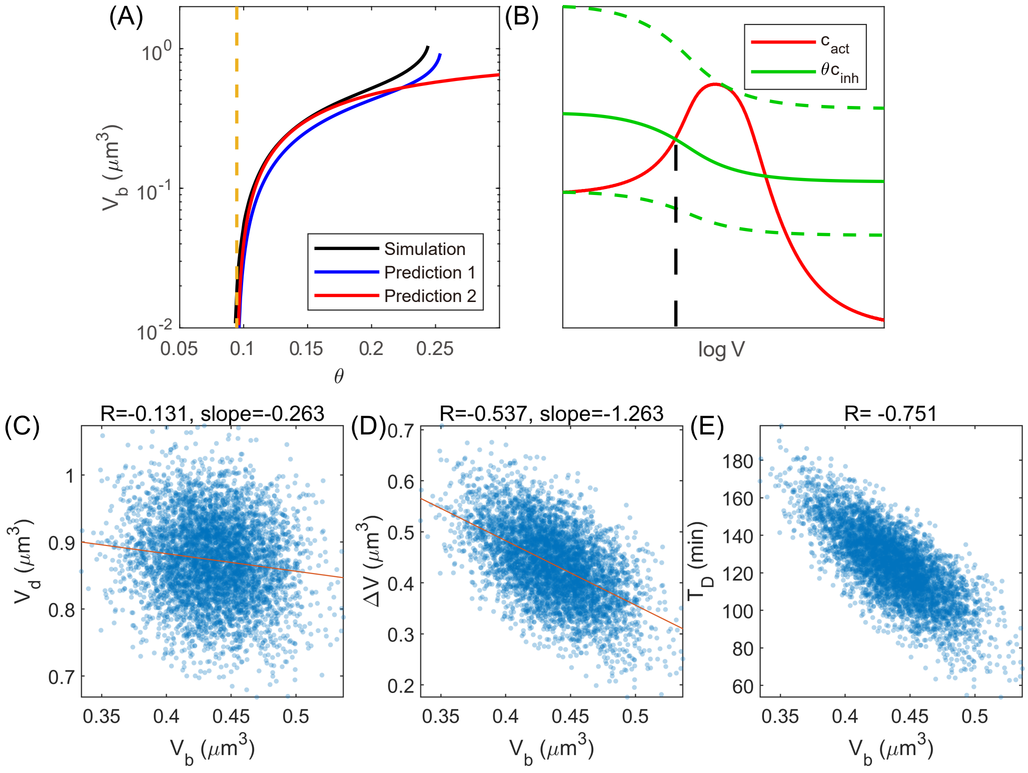
(A)
(B)
(C)
(D)
(E)
Simulation
Prediction 1
Prediction 2
$V_b$ ($\mu$m$^3$)
$\theta$
$c_{act}$
$\theta c_{inh}$
log V
R=-0.131, slope=-0.263
R=-0.537, slope=-1.263
R= -0.751
$V_d$ ($\mu$m$^3$)
$\Delta V$ ($\mu$m$^3$)
$T_D$ (min)

S7 Fig.

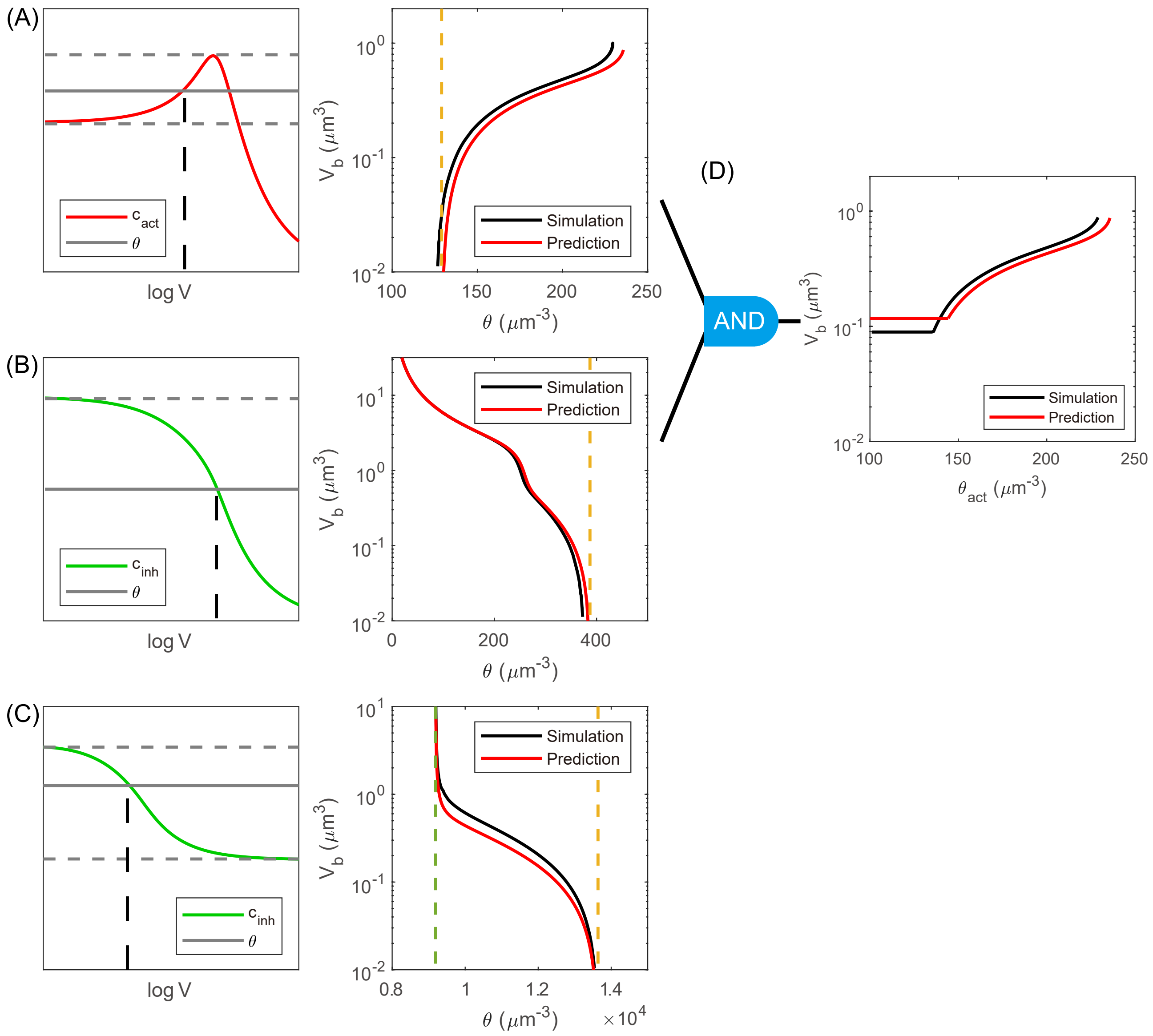

(A)
c act
θ
log V
10^0
10^-1
10^-2
V b (μm^3)
100
150
200
250
θ (μm^-3)
Simulation
Prediction
(B)
c inh
θ
log V
10^1
10^0
10^-1
10^-2
0
200
400
(C)
10^1
0.8
1
1.2
1.4
×10^4
(D)
AND
θ act (μm^-3)


S8 Fig.

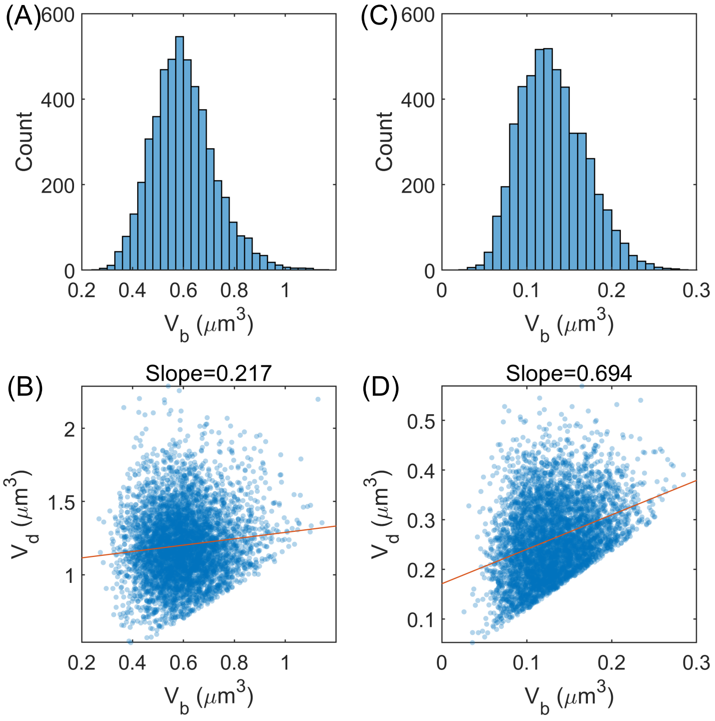
(A)
600
400
200
0
Count
0.2
0.4
0.6
0.8
1
$V_b$ ($\mu m^3$)
(C)
0
0.1
0.2
0.3
(B)
Slope=0.217
2
1.5
1
$V_d$ ($\mu m^3$)
(D)
Slope=0.694
0.5
0.4
0.3
0.2
0.1

S9 Fig.

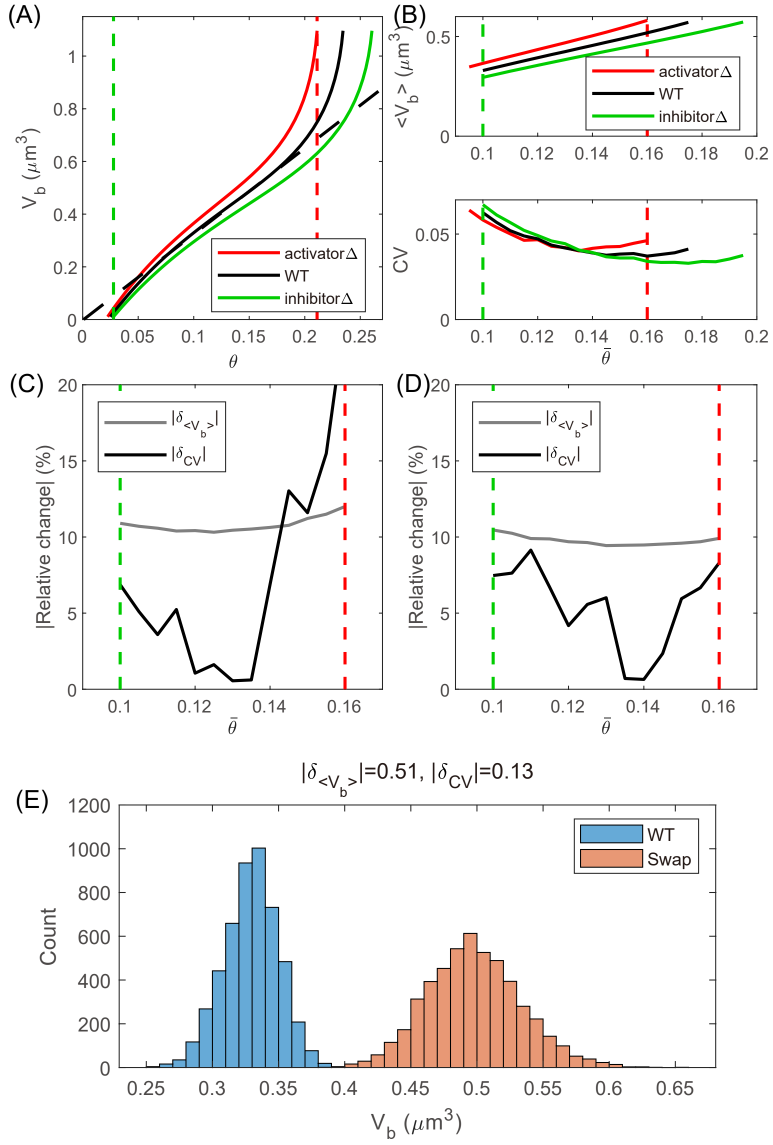
(A)
(B)
(C)
(D)
(E)
activatorΔ
WT
inhibitorΔ
$V_b$ ($\mu m^3$)
$\theta$
$<V_b>$ ($\mu m^3$)
CV
$\bar{\theta}$
|Relative change| (%)
$|\delta_{<V_b>}|$
$|\delta_{CV}|$
$|\delta_{<V_b>}|$=0.51, $|\delta_{CV}|$=0.13
WT
Swap
Count

S10 Fig.

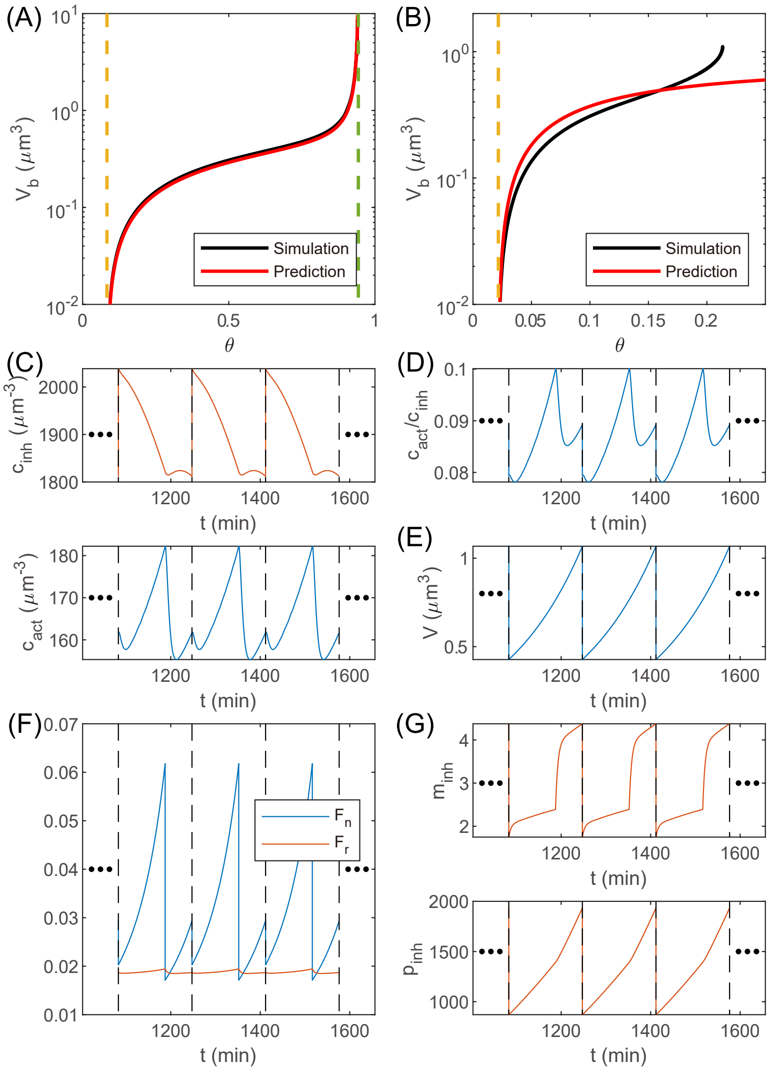

(A)
(B)
Simulation
Prediction
$V_b$ ($\mu m^3$)
$\theta$
(C)
(D)
(E)
(F)
(G)
$c_{inh}$ ($\mu m^{-3}$)
$c_{act}/c_{inh}$
$c_{act}$ ($\mu m^{-3}$)
V ($\mu m^3$)
$F_n$
$F_r$
$m_{inh}$
$p_{inh}$
t (min)


S11 Fig.

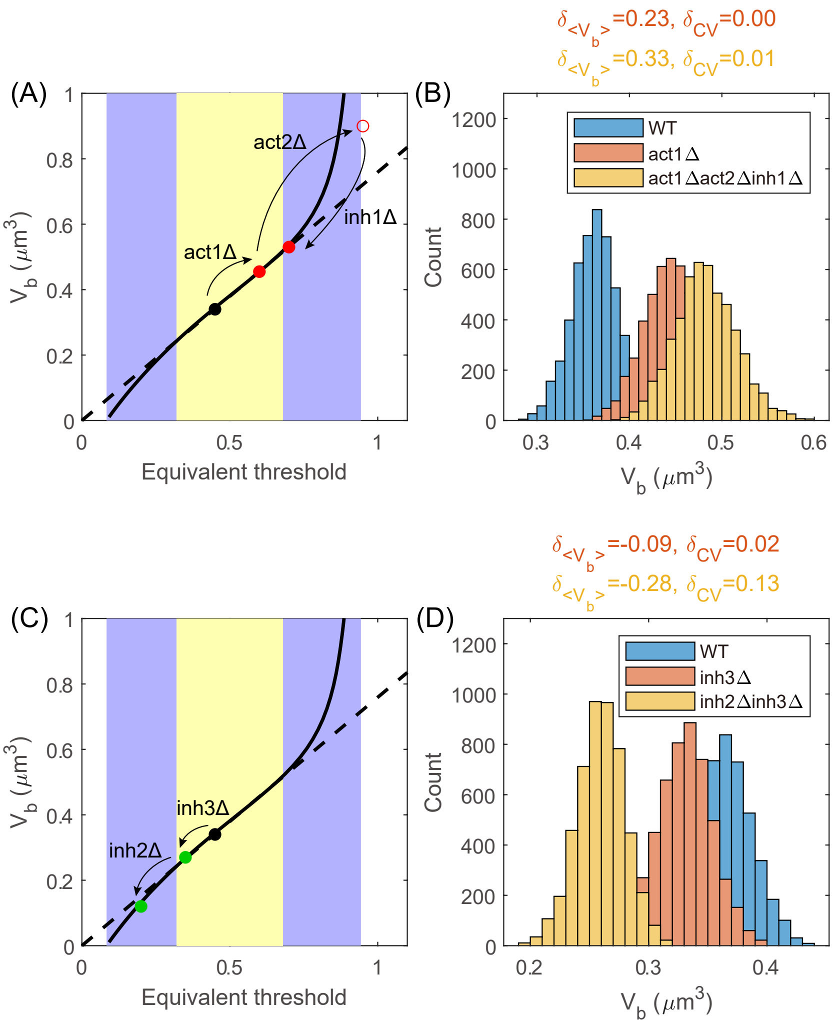

δ<Vb>=0.23, δCV=0.00
δ<Vb>=0.33, δCV=0.01
(A)
(B)
act2Δ
act1Δ
inh1Δ
Vb (μm3)
Equivalent threshold
WT
act1Δ
act1Δact2Δinh1Δ
Count
Vb (μm3)
δ<Vb>=-0.09, δCV=0.02
δ<Vb>=-0.28, δCV=0.13
(C)
(D)
inh3Δ
inh2Δ
WT
inh3Δ
inh2Δinh3Δ


S12 Fig.
